# Dealing with the promise of metabarcoding in mega-event biomonitoring: EXPO2015 unedited data

**DOI:** 10.1101/2022.01.02.474438

**Authors:** Giulia Agostinetto, Antonia Bruno, Anna Sandionigi, Alberto Brusati, Caterina Manzari, Alice Chiodi, Eleonora Siani, Luigimaria Borruso, Andrea Galimberti, Graziano Pesole, Massimo Labra, Maurizio Casiraghi

## Abstract

As human activities on our planet persist, causing widespread and irreversible environmental degradation, the need to biomonitor ecosystems has never been more pressing. These circumstances have required a renewal in monitoring techniques, encouraged by necessity to develop more rapid and accurate tools which will support timely observations of ecosystem structure and function. The World Exposition (from now ‘EXPO2015’) hosted in Milan from May to October 2015 was a global event that could be categorized as a mega-event, which can be defined as an acute environmental stressor, possibly generating biodiversity alteration and disturbance.

During the six months of EXPO2015, exhibitors from more than 135 countries and 22 million visitors insisted on a 1.1 million square meters area. Faced with such a massive event, we explore the potential of DNA metabarcoding using three molecular markers to improve the understanding of anthropogenic impacts in the area, both considering air and water monitoring. Furthermore, we explore the effectiveness of the taxonomy assignment phase considering different taxonomic levels of analysis and the use of data mining approaches to predict sample origin. Unless the degree of taxa identification still remains open, our results showed that DNA metabarcoding is a powerful genomic-based tool to monitor biodiversity at the microscale, allowing us to capture exact fingerprints of specific event sites and to explore in a comprehensive manner the eukaryotic community alteration. With this work, we aim to disentangle and overcome the crucial issues related to the generalization of DNA metabarcoding in order to support future applications.

## Introduction

Environmental degradation due to anthropic activities have increased the scale and frequency of biodiversity assessments. Certainly, the environmental degradation is particularly dramatic in the highly anthropogenic areas. Governments and international organisations are issuing and thus including in their agendas new strategies to protect and restore biodiversity, such as the Intergovernmental Science-Policy Platform on Biodiversity and Ecosystem Services (IPBES 2019; Bongaarts, 2019; Lanzen et al., 2016; Baird et al., 2012). These circumstances have required a renovation in monitoring techniques, encouraged by the necessity to develop more rapid and accurate tools supporting timely observations of ecosystems structure and functions (Taylor et al., 2016; Pimm et al., 2015). In this framework, supported by Next Generation Biomonitoring (NGB) initiatives (Makiola et al., 2020), DNA metabarcoding introduced surprising signs of progress in surveying prokaryotic and eukaryotic diversity from any type of environment (Makiola et al., 2020; McGee et al., 2019). After a few years from the adoption of DNA metabarcoding, many worldwide molecular data collection projects include DNA metabarcoding data, as a natural progression for biodiversity assessment e.g. Earth BioGenome Project (EBP 2021), The European Reference Genome Atlas initiative (ERGA 2021), the BIOSCAN project (BIOSCAN2021), the Vertebrate Genomes Project (VGP) (Rhie et al. 2021), the i5k Arthropod Genomes Initiative (i5K Consortium 2013), the 10KP Plant Genomes Project (Cheng et al. 2018), and others (Waterhouse et al 2021). DNA metabarcoding widespread adoption has also been supported by the advances in high-throughput DNA sequencing (HTS) technologies, increasing data yield with costs reduction (Cordier et al., 2020; Porter et al., 2018; Pimm et al., 2015; Thomsen et al., 2015; Shokralla et al., 2012), allowing taxa exploration at unprecedented extent, for a time and cost-effective biodiversity monitoring (Westfall et al., 2019; Ruppert et al., 2019; Deiner et al., 2017).

Several studies exploited the potential of DNA metabarcoding to improve the understanding of anthropogenic impacts (Cordier et al., 2021; Tommasi et al., 2021; Frontalini et al., 2018; Lanzen et al., 2016), monitoring alien species introduction (e.g., Westfall et al., 2019; Comtet et al., 2015), even in the context of regulatory policymaking (Cordier et al., 2021; van der Heyde et al. 2020; Pawlowski et al., 2018).

However, the DNA metabarcoding data analysis and interpretation still requires great efforts to achieve full data exploitation and not standard procedures could be applied for each taxa domain (Ruppert et al., 2019; Porter et al., 2018; Deiner et al., 2017). In particular, difficulties remain related to the lack of information in reference taxonomic databases (Curry et al., 2018; Weigand et al., 2019), taxonomic resolution and misidentification (Bush et al., 2019), leading also to the implementation of taxonomy free approaches (Vasselon et al., 2017).

Nevertheless, we harnessed the advantages and the huge amount of information that DNA metabarcoding can generate to investigate the influence of massive human-induced activities on biological communities, also considering the issues related to marker choice, reads processing and the information contained in reference databases, a fundamental part of data interpretation.

In this study, we focused on the monitoring of the World Exposition (from now “EXPO2015”) hosted in Milan from May to October 2015. This global event was categorized as a mega-event (Muller, 2015), which can be defined as an acute environmental stressor, possibly generating biodiversity alteration and disturbance.

During the six months of EXPO2015, exhibitors from more than 135 countries and 22 million visitors insisted on a 1.1 million square meters exhibition area. Faced with such a massive event, a wide-range analysis of biodiversity could be reliable for addressing biomonitoring purposes (Cordier et al., 2021; Cristescu et al., 2018; Alberdi et al., 2018; Trebitz et al., 2017; Comtet et al. 2015). To overcome restrictions of traditional biomonitoring, which is limited to observations on small sets of bioindicators and/or flagship species (Cordier et al., 2021; Dequiedt et al. 2011; Magurran et al. 2010; Reavie et al. 2010; Bonada et al. 2006), we applied a DNA metabarcoding approach targeting three different molecular markers and involving two different sampling strategies (i.e., water and air) to obtain a comprehensive overview of the impact of the exhibition on environmental community assemblages. In this context, both overall and microscale investigations were conducted. Specifically, we monitored the water canalization system, which connects two local rivers across the exhibition area, the two local rivers and the air biodiversity collected at two representative sites. We chose three mini-barcode regions allowing the assessment of a broad taxonomic spectrum of the eukaryotic community: the V9 hypervariable region of 18S SSU rRNA (Harrison et al., 2021; Fernández-Álvarez et al. 2018; Chariton et al. 2015; Cowart et al. 2015; Lallias et al. 2015; Zimmermann et al. 2015; Edgcomb et al., 2011), the plastid trnL intron (Deiner et al., 2017; Fahner et al., 2016; Quéméré et al., 2013; Taberlet et al., 1991), and the internal transcribed spacer ITS2 of rRNA (Nilsson et al., 2019; Blaalid et al., 2013; Toju et al., 2012; White et al., 1990).

Overall, our main intent was to validate DNA metabarcoding as a biomonitoring strategy to understand the environmental impact of global events, such as EXPO2015, on eukaryotic community diversity. In parallel, we tried to deepen the following questions: i) if DNA metabarcoding can track biodiversity communities in a mega-event context, ii) which are the pros and cons of using multi-marker strategies, considering the absence of common procedures and the issues related to the taxonomy assignment and iii) if machine learning strategies can help in predicting sample origin, overcoming the uncertainty of the taxonomy assignment.

Our results showed that DNA metabarcoding coupled with machine learning approach is a powerful genomic-based tool to monitor biodiversity at the microscale, allowing us to capture exact fingerprints of specific event sites and to explore in a comprehensive manner the eukaryotic community alteration. We discussed in the work the crucial issues related to the generalization of the approach and the degree of taxa identification. We provided a case-study application of DNA metabarcoding to an urban context, monitoring biodiversity at micro-scale, but also with a focus on the changes starting from the laying of the first stone. As well as the great potential of genomic-based tools and data related to genetic biodiversity are growing, machine learning approaches could give the decisive breakthrough to the application of DNA-based monitoring 3.0 at a broader extent.

## Materials and methods

### Study area and sampling

The EXPO 2015 exhibition site is located northwest of Milan. The site occupies an area of 110 hectares, with approximately 250,000 m^2^ of vegetation, 6,000 m of canals, more than 70 exposition pavilions, for the exhibitors coming from more than 135 countries, built in three and a half years, and was completed just hours before the opening ceremony (Expo Milano 2015 Official Report ^1^).

It had long been an industrial zone before its conversion to logistical and municipal services and agriculture. The area is characterized by two parallel water canals and it is crossed by two rivers, Guisa and Olona.

Within the EXPO area, four main sampling points were considered (Fig. 1):

**Figure 1.**
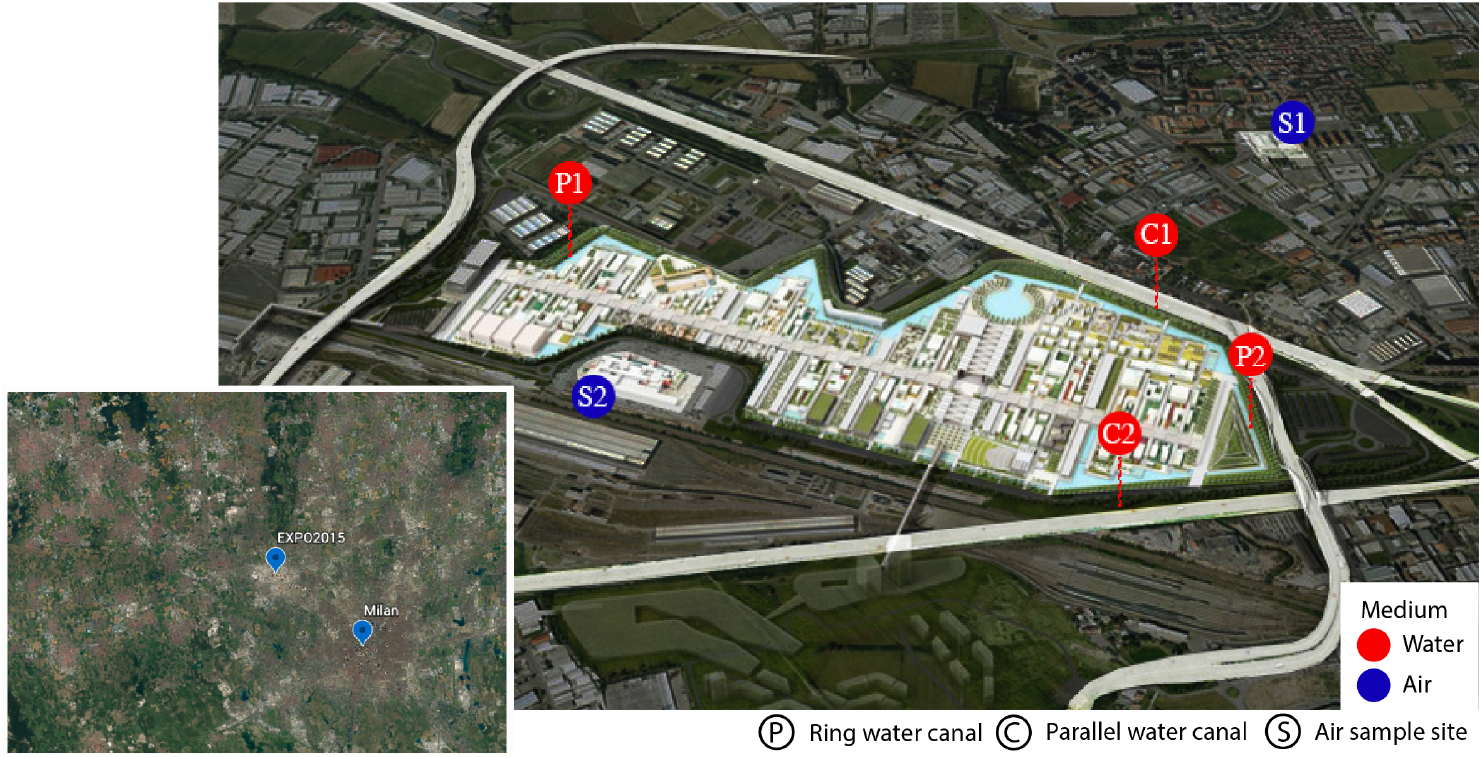
Map of sampling sites. Blue circles indicate the two sampling points of air (S2 closer to the exposition site). Red circles indicate the four sampling points of water canals (P1 P2 ring water canal; C1 C2 water canal parallel).

- P1: localized in the ring water canal, upstream of P2 (inlet water);
- C1: located along the water canal parallel to the area, receives incoming water from the Guisa river and enters more times in contact with the area;
- P2: localized in the ring water canal, downstream of P1, collects outlet water derived from the whole area and from P1;
- C2: located along the water canal parallel to the area, receives the water from the exhibition area and enters the river Olona.

Considering the water sampling, samples were collected using one-liter sterile single-use bottles (LP Italia) in PET from the two rivers crossing the area of the exhibition, Guisa and Olona, and at four sites localized within the EXPO area (P1, P2, C1 and C2; see Figure 1). Sampling began in October 2014 and ended in March 2016. Since the works for the construction of the exhibition site have continued up to the days immediately prior to the opening of the event, the sampling of water perimeter channels was not possible in the *ante operam* phase (i.e., before May 2015) in the EXPO area.

Regarding the *post operam* phase (i.e., after October 2015), the analyzed samples were collected at the same sampling sites, since the exposition area was no longer accessible. In total, Guisa was sampled six times (6 samples) and Olona three times (3 samples) as the river was dry, once a month (for details about sampling dates, see Supplementary S1).

The sites C1, C2, P1, and P2 were sampled monthly during the EXPO event (*in operam* phase), obtaining 30 samples of P1, 18 for C1, 33 for P2 and 26 for C2.

Considering the air sampling campaign, samples were collected monthly from October 2014 to March 2016 through two different methods: a Tauber Trap approach (Tauber, 1974) and Lanzoni VPPS 2010 (**Lanzoni**, Bologna, Italy) instrument (based on Hirst model; Hirst, 1952; Núñez et al., 2017)

Sites sampled were:

- S1, located at the company Tarenzi s.p.a, 600 meters north of EXPO (a total 44 samples);
- S2, located on the roof of c.m.p. Poste Roserio, 100 meters south of EXPO (a total of 47);

The S1 site was investigated using the Tauber Trap method, instead of the S2 site in which both instruments were installed. Sites were carefully selected for their geographical position, near the exhibition area and opposite each other, in order to collect the biological component considering wind direction. The different distance of sampling sites from the EXPO area allowed both short-range (100 meters) and long-range (600 meters) monitoring (c.m.p Roserio and Tarenzi s.p.a., respectively). Overall, a total of 228 samples were collected from water (137 samples) and air (91 samples), covering the period from October 2014 to March 2016 (for time point list see Tables in Supplementary S1), using three molecular markers. The sample distribution was conducted as follows: 34 air and 47 water samples belonging to 18S V9 region, 30 air and 45 water samples to trnL and 27 air and 45 water samples belonging to ITS2.

### Samples pre-processing and environmental DNA extraction

Each liter of water belonging to each site was pre-processed with serial filtrations with the use of nitrocellulose and acetate membrane filters with 8 μm and 0.45 μm pore sizes (Jamwal et al., 2021; Valsechi et al., 2021; Capo et al., 2020), respectively. For the air sampling campaign, each Tauber trap sample (composed by a solution of ethanol and glycerol) was pre-processed with serial filtrations with the same strategy used for water samples.

Filters belonging to both media were initially crushed with Tissue-Lyser and liquid nitrogen. Subsequently, the DNA was extracted using the EuroGold Plant DNA Mini Kit (EuroClone). DNA extraction from samples subjected to mechanical lysis was carried out following the protocol for dry material with the following modifications: instead of starting from 250 mg of dry material, all the filters obtained for each sample were processed together, so that the DNA extracted corresponded to the volume of filtered water. DNA elution was carried out with 100 μl of elution buffer.

Three genetic markers (i.e, the nuclear V9 region of 18S rDNA and ITS2 and the plastid intron trnL) have been selected. The V9 region of 18S rDNA was used as a generalist genetic marker to explore the eukaryotic community (Harrison et al., 2021; Fernández-Álvarez et al. 2018; Chariton et al. 2015; Cowart et al. 2015; Lallias et al. 2015; Zimmermann et al. 2015; Edgcomb et al., 2011). The plastid intron trnL and the internal transcribed spacer ITS2 were used specifically to identify Plantae (Deiner et al., 2017; Fahner et al., 2016; Quéméré et al., 2013; Taberlet et al., 1991) and Fungi (Nilsson et al., 2019; Blaalid et al., 2013; Toju et al., 2012; White et al., 1990), respectively.

Raw reads were generated in an eighteen month assessment (from October 2014 to March 2016; Supplementary Files S1), collecting a total of 228 samples (i.e., 137 water and 91 air), sequenced at the three selected loci markers (Supplementary Table S3).

### Illumina library preparation and sequencing

V9 hypervariable region of 18S rRNA gene, intron trnL and ITS2 (primer details are provided in Supplementary Table S3) libraries were generated following the standard protocol (16S Metagenomic Sequencing Library Preparation, Part # 15044223 Rev. B). Amplicon PCRs were performed using the primer pairs used for qPCR quantification plus the adapter sequence. Libraries were quantified with a 2100 Bioanalyzer (Agilent Technologies) and sequenced with the Illumina MiSeq platform (five runs, v2 chemistry, 2×150bp). Library preparation and sequencing were carried out at IBIOM-CNR (Bari, Italy). Quantification protocol and primer list are available in Supplementary Data S2.

### Bioinformatics workflow, biodiversity and machine learning analysis

For each marker gene, the raw paired-end FASTQ reads were imported into the Quantitative Insights Into Microbial Ecology 2 program (QIIME2, ver. 2020.8; Bolyen et al., 2019) and demultiplexing native plugin. Illumina runs were processed independently with the Divisive Amplicon Denoising Algorithm 2 (DADA2) plugin (Callahan et al., 2016). DADA2 was used to filter, trim, denoise, merge, remove of chimeras and calculate ESVs (Exact Sequence Variants; Callahan et al. 2017). In particular, an expected error = 2.0 was used as an indicator of read accuracy. Primers were trimmed and low-quality bases were removed. ESVs sequences with at least 10 representatives were taxonomically assigned using OBITools (Boyer et al., 2015) by ecotag tool, comparing sequences with an ecoPCR database extracted from the EMBL database version r139 (Kanz et al., 2004).

For each marker gene, the results of the taxonomy assignment were analysed considering the percentage of rank assigned at different levels (Kingdom, Phylum, Class, Order, Family, Genus, Species).

In order to estimate the biodiversity variation, we calculated alpha and beta diversity index for each marker gene separately. In detail, differences among sample types (water and air), sites (sampling points) and the macro category (air: S; rivers: R; internal canals: P; external canals: C) were tested For alpha diversity, we considered Shannon metric and presence/absence observations. Differences were tested using the pairwise Krustall-Wallis test implemented in the alpha-group-significance QIIME2 plugin (Kruskal and Wallis, 1952). To assess how volatile a dependent variable (alpha diversity measured as Shannon diversity) is over an independent variable (time) in water and air medium, a volatility plot was generated for each marker. For beta diversity, we calculated Jaccard metric to test differences among sample types, sites and macro categories using a PERMANOVA analysis performed with beta-group-significance plugin (Anderson, 2001).

Subsequently, the Random Forest classifier implemented in the sample-classifier QIIME2 plugin (Bokulich et al., 2018) was used to classify samples based on sites and macro categories metadata. The number of trees to grow for estimation was set to 1,000. Overall accuracy (i.e., the fraction of times that the tested samples are assigned the correct class) was calculated for each factor. K-fold cross-validation was performed during automatic feature selection and parameter optimization steps. A fivefold cross-validation was also performed. Further, machine learning analysis was carried out considering the genetic information of all the three marker regions, based on sites and macro categories metadata.

Figures and plots were created through QIIME2 plugins (Bolyen et al., 2019; Anderson, 2001; https://github.com/qiime2/q2-taxa) and ExTaxsI tool (Agostinetto et al., 2021; Agostinetto et al., 2020; https://github.com/qLSLab/ExTaxsI) to give an overview of biodiversity collected during the sampling campaign, with the aim to summarize the great amount of data generated and help data interpretation. Raw reads were submitted to the ENA database and they will be made public upon paper acceptance.

## Results

### Sequencing results

Nine Illumina MiSeq sequencing runs for the three markers selected (18S SSU rRNA, trnL and ITS2) produced a total of 127,971,220 reads (63,985,610 pair-end reads), belonging to 228 samples. After the filtering steps, a total of 44,193,721 sequences were retained for the downstream analysis. As the DADA2 R package implements a full amplicon workflow (Callahan et al., 2016), we obtained a total of 19,304 ESVs (Callahan et al., 2017) for V9 raw reads, 3,630 ESVs for trnL and 8,471 ESVs for ITS2. Complete ESVs tables are available in Supplementary S8.

### Taxonomy results

V9 18S sequences resulted in 77.43 % assigned to Unicellular Eukaryotes, 13.20% to Fungi, 6.9% to Viridiplantae group, 3.58% to Metazoa and 2.54% to Bacteria, with 20.01% Unassigned sequences. Overall, 44.28% of assignments reached the genus level (Figure 5). Among Metazoa assignments, 39.65 % was composed by Arthropoda, 13.60% by Nematoda and 9.26 % by Rotifera. These taxa were followed by Platyhelminthes (7.8%), Unknown sequences (6.80%), Gastrotricha (6.80%), Annelida (4.62%), Cnidaria (3.76%), Chordata (2.31%), and Tardigrada (2.02%). A small fraction of assignments collected Mollusca (1.44%), Porifera (1.15%), Bryozoa (0.28%), Ctenophora (0.28%) and Nemertea (0.14%). Among Metazoa sequences, 45.5% of them reached a genus level assignment.

ITS2 sequences resulted in 64.29% of Fungi assignments, followed by 17.99% of Unclassified Eukaryotes, 11% of Unassigned sequences, 8% of Viridiplantae and 0.36% of Metazoa sequences. Overall, 46.86% of sequences reached a genus level assignment. Among Fungi sequences, 29.99% of them were assigned to Ascomycota phylum and 28.54% to Basidiomycota phylum, followed by 2.23% of Chytridiomycota, 0.56% of Mucoromycota, 0.04 % of Zoopagomycota, 0.01% of Olpidiomycota and 0.01% of Blastocladiomycota.

Plastid trnL intron sequences resulted in 51.81% of Streptophyta assignments, followed by 42.17% of Viridiplantae Unassigned sequences and 6% of Chlorophyta sequences. Overall, 14.38% of sequences reached the genus level assignment. Among Streptophyta, 63.42% remained Unassigned.

For each marker gene, the distribution of taxa among sites can be consulted into the respective resume figures, in particular considering Metazoa for V9 18S, Fungi for ITS2 and Streptophyta for trnL (Figure 2-3-4, section “c”). In addition, tables with the complete taxonomy assignments are available in Supplementary S9.

**Figure 2.**
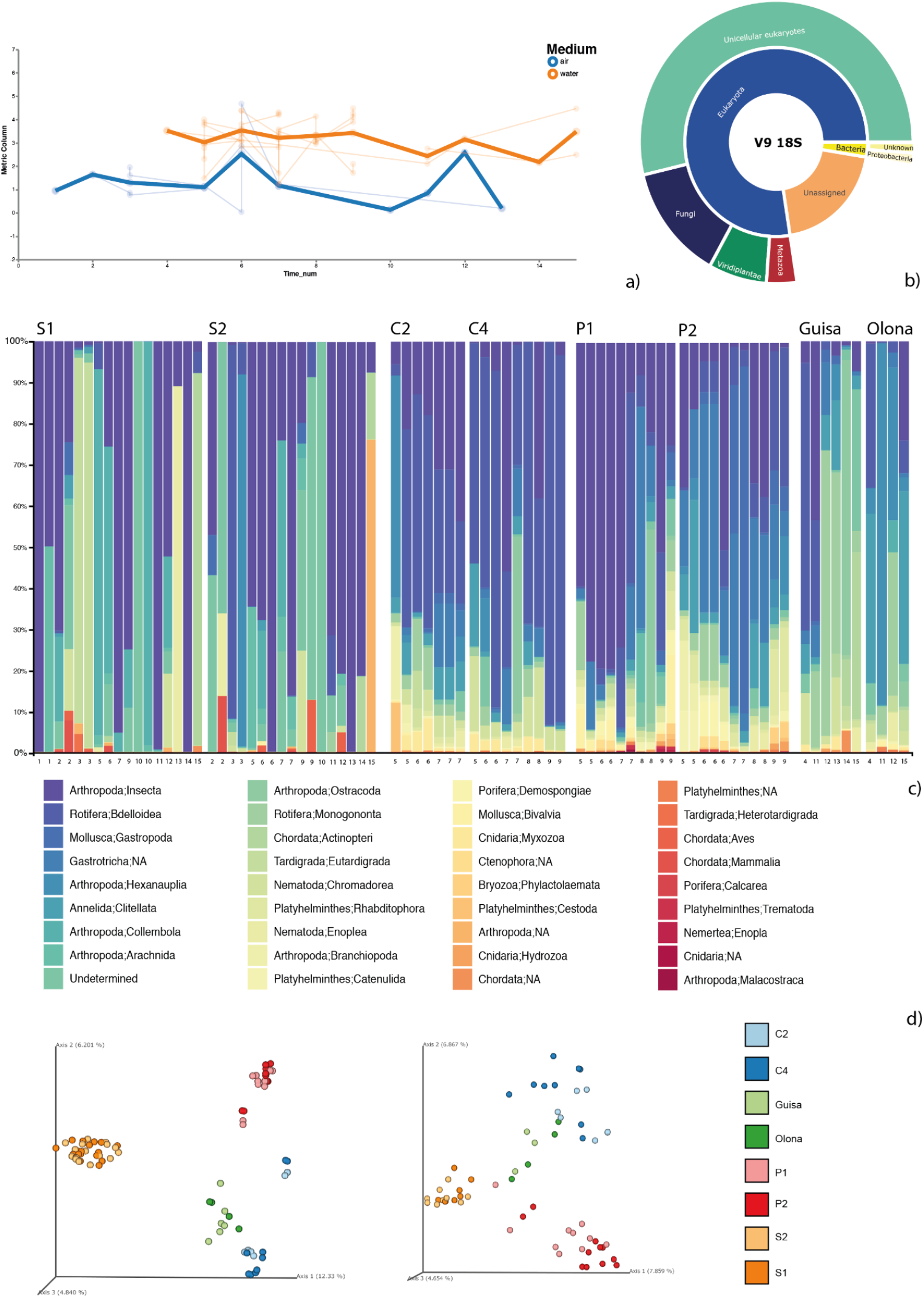
Resume figure showing the V9 18S results with: a) volatility plot and b) sunburst plot representing the taxa distribution of taxonomy assignment; c) taxa-bar-plot considering only Metazoa assignments; d) PCoA analysis based on Jaccard metric considering ESVs (left) and ESVs assigned only to Metazoa (right) on sampling sites.

**Figure 3.**
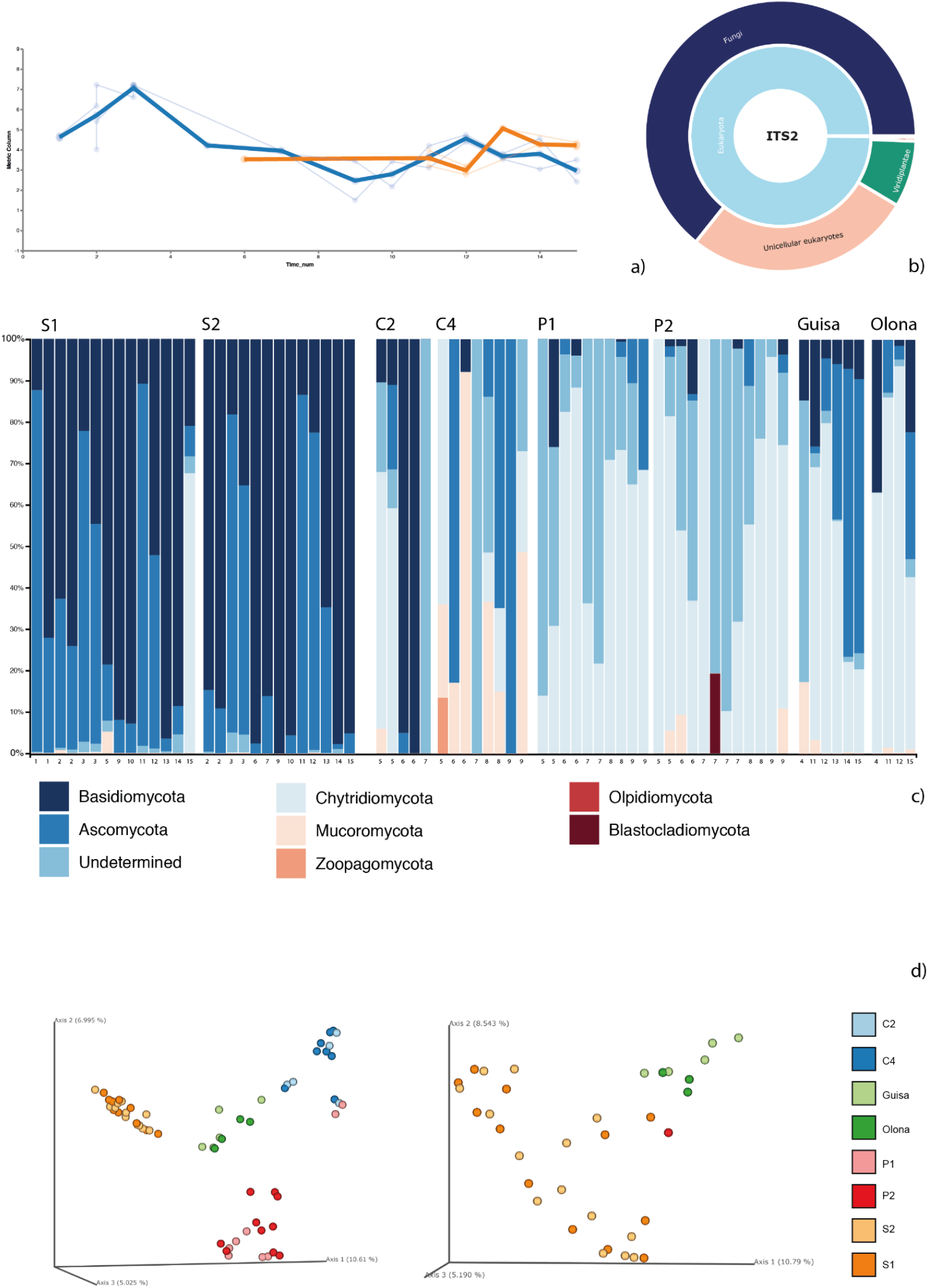
Resume figure showing the ITS2 results with: a) volatility plot and b) sunburst plot representing the taxa distribution of taxonomy assignment; c) taxa-bar-plot considering only Fungi assignments; d) PCoA analysis based on Jaccard metric considering ESVs (left) and ESVs assigned only to Fungi (right) on sampling sites.

**Figure 4.**
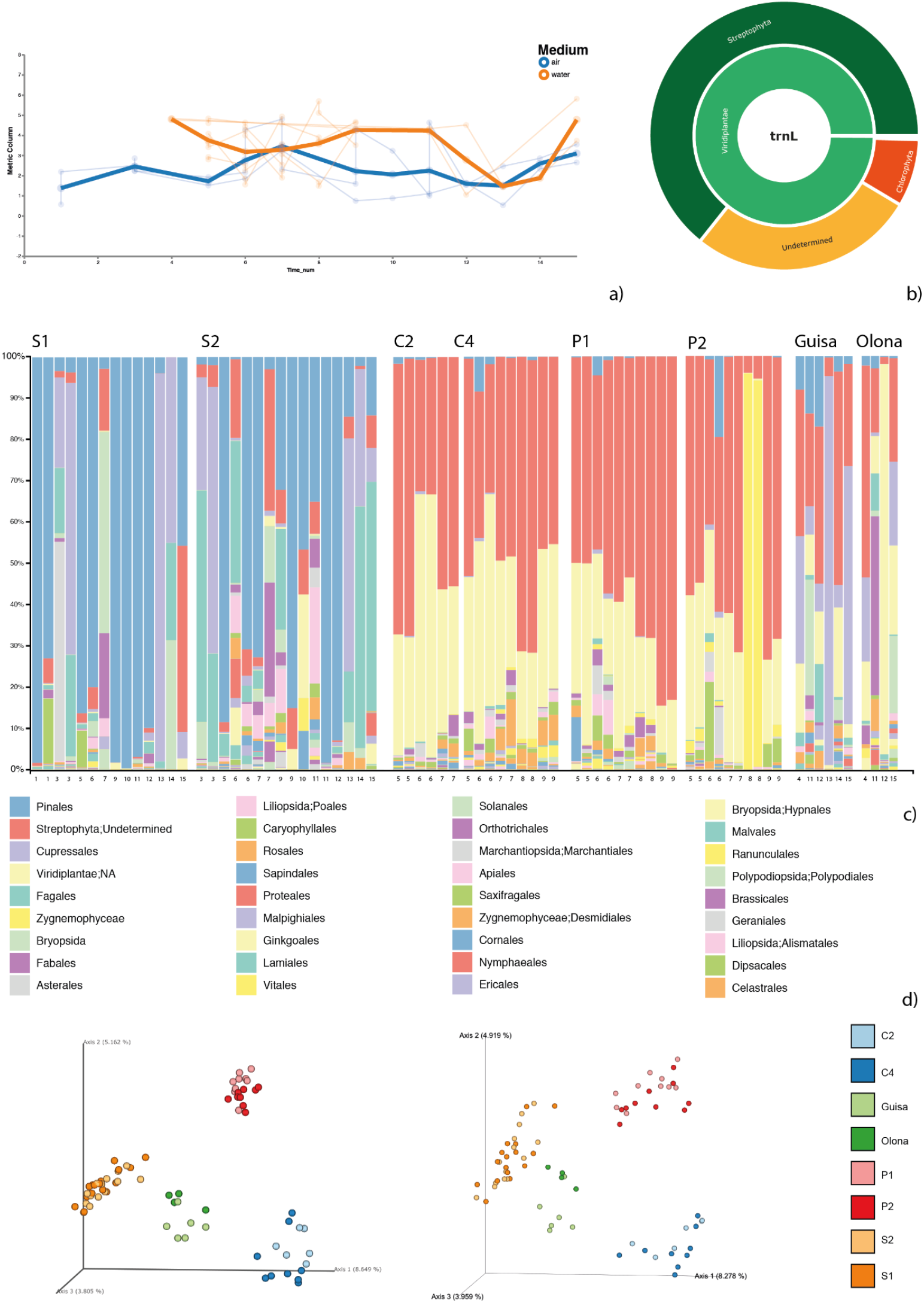
Resume figure showing the trnL results with: a) volatility plot and b) sunburst plot representing the taxa distribution of taxonomy assignment; c) taxa-bar-plot considering only Streptophyta assignments; d) PCoA analysis based on Jaccard metric considering ESVs (left) and ESVs assigned only to Streptophyta (right) on sampling sites.

To explore the results of the taxonomic assignment for each rank, we provide in Figure 5 a report with the percentage of sequences assigned for each marker region (complete data are available in Supplementary S4). In detail, we explored for each marker gene the taxa reached during the taxonomy assignment. The trnL intron was the only marker for which we assigned the 100% of the sequences obtained, in particular to Viridiplantae Kingdom. In addition, at least the 10% of ESVs with a complete Phylum and Order rank lacked Class information. Considering the number of species assigned, trnL was the marker which performed the worst.

**Figure 5.**
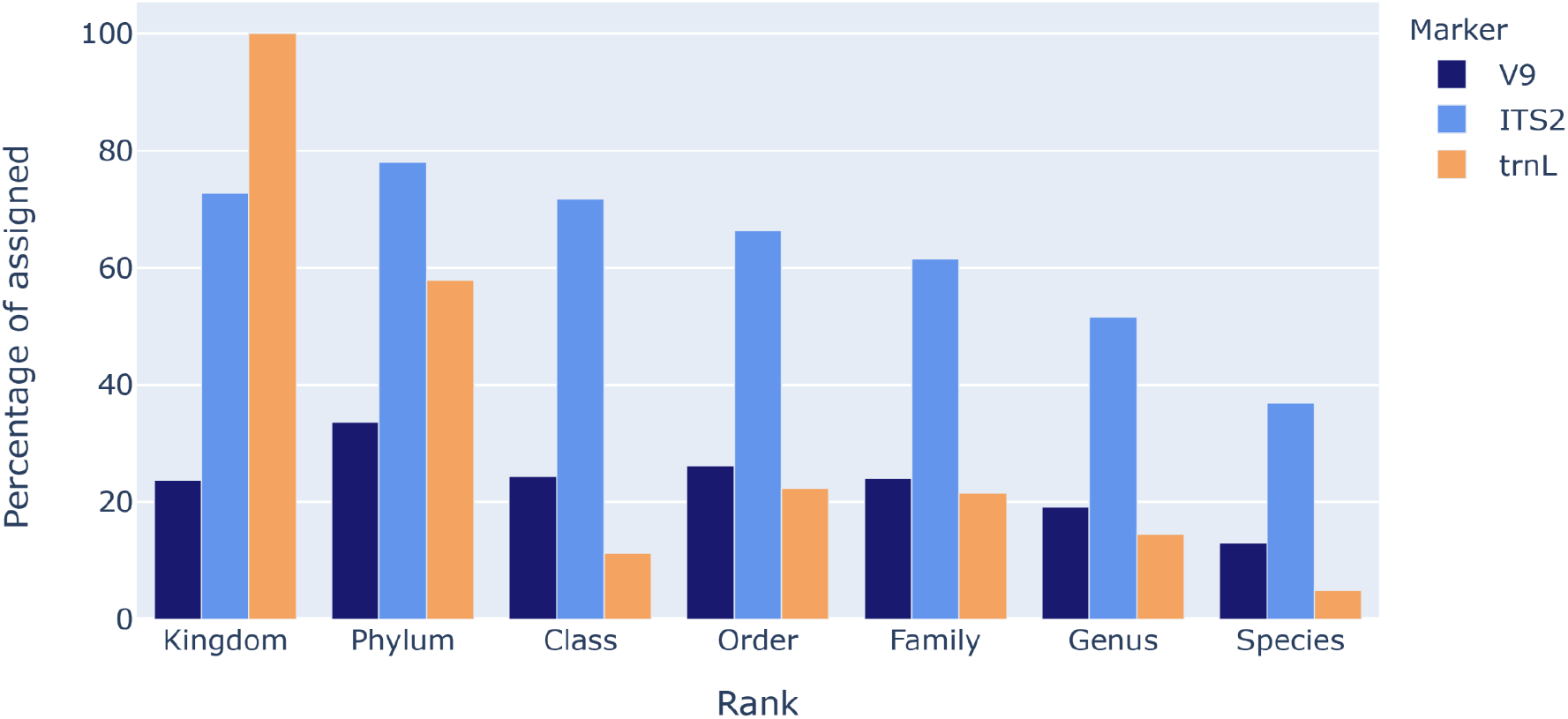
Percentage of sequences assigned for each marker region considering a 7 rank NCBI taxonomy (complete data are available in Supplementary S4).

From ITS2, we obtained the best results considering the taxonomy assignment, since the number of assigned ranks gradually decreased without gaps between one rank and another. Issues were found considering the Kingdom level: the Kingdom rank was not specified for taxa belonging to the Unicellular Eukaryotes group and a lower percentage of sequences were assigned if we consider the Kingdom rank instead of the Phylum.

V9 18S had the same issues of ITS2 considering Kingdom and Phylum ranks, as sequences belonging to V9 18S were also assigned to the Unicellular Eukaryotes group. For this reason, issues related to Class information were observed. In general, it was the marker with the highest percentage of Unassigned sequences.

A summarization of assignments was represented with sunburst plots in Figure 2-3-4 section “b”, accompanied by the results related to the ranks assigned in Figure 5. In addition, for each marker we added interactive sunburst charts obtained via ExTaxsI tool to explore dynamically the taxonomy obtained (Supplementary S7; Agostinetto et al., 2021; Agostinetto et al., 2020).

### Biodiversity analysis

For each genetic marker, a biodiversity analysis was performed considering the type of sample, the sampling site and the macro category (air: S; rivers: R; internal canals: P; external canals: C).

The data analysed consisted not only of the ESVs calculated with the bioinformatics analysis, but also filtering the assigned ESVs considering different taxonomic levels, based on the results of the taxonomy assignments. In particular, for V9 18S we considered sequences assigned only to the Metazoa group and also sequences assigned to eukaryotic taxa, overall. For ITS2, sequences assigned to Fungi were considered. For trnL, Streptophyta sequences and sequences excluding Chlorophyta were considered. Finally, A PERMANOVA test was used to assess statistical significance.

Considering alpha diversity analysis, a significant difference was observed between air and water samples of V9 18S and trnL. In addition, considering V9 18S, a difference was found between macro category samples of each group considered, in particular between C and P. For ITS2 sequences a difference was detected among macro category sites (C-P and C-R). A stronger difference was detected considering Fungi sequences, also regarding water samples sites (C2-GU; GU-P1; GU-P2). For trnL, all the three types of analysis (only ESVs, Streptophyta sequences and sequences excluding Chlorophyta) showed a difference among water and air samples. For details about alpha diversity results, see Supplementary S5a.

Considering beta diversity, sample medium (air and water), sampling sites and macro category were analysed. A significant difference was observed between the two different sampling media, for all the markers tested, both considering ESVs and taxa detected. In addition, significant differences were found comparing both macro category and sampling sites, both considering ESVs and taxa detected. For details about PERMANOVA results, see Supplementary Material S5b.

Overall, PCoA plots (Figure 2-3-4, section “d”) showed a significant structuration (model results are reported in Supplementary Materials S5) based on sampling site (different sampling point in EXPO2015 area), with the Internal sites (P1-P2) clustering close to each other, as well as the External sites (C1-C2) and Rivers (OL-GU). The same significant structure is also visible considering the taxonomic information, for which we reported in the main figures the results about Metazoa group (18S V9), Fungi (ITS2) and Streptophyta (trnL) (Figure 2-3-4 section “d”).

### Machine learning analysis

DNA metabarcoding monitoring studies often aim to differentiate samples based on their biodiversity composition, a task that can be efficiently performed by Supervised Learning methods (Knights et al., 2011, Bokulich et al, 2018). We used a supervised machine learning approach to evaluate the potential of DNA metabarcoding data to classify sampling sites and macro category outcomes, considering the different types of information that we obtained based on the taxonomy assignments results (see the section above).

For all the three markers analysed, an improvement of classification was seen passing from sites to macro category metadata prediction. This trend is visible in Figure 6 for V9 18S and in Supplementary S10 for trnL and IT2.

**Figure 6.**
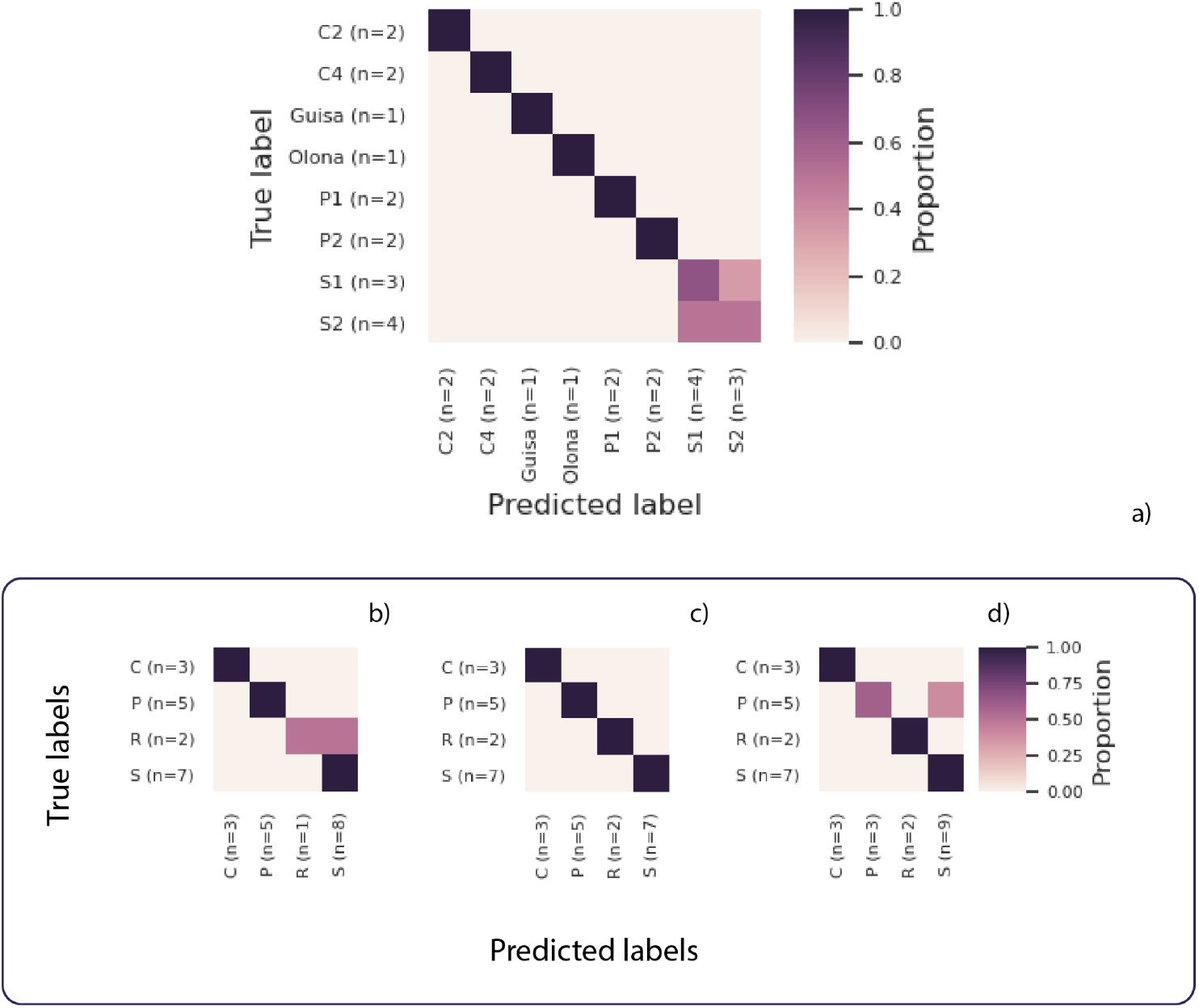
SL results of V9 18S marker: results were reported considering a) sequences per sites and considering the division of sites into macro categories of b) sequences, c) Eukaryota, d) Metazoa. The Figure shows a scatter plot of true vs. predicted values for regression results.

**Figure 7.**
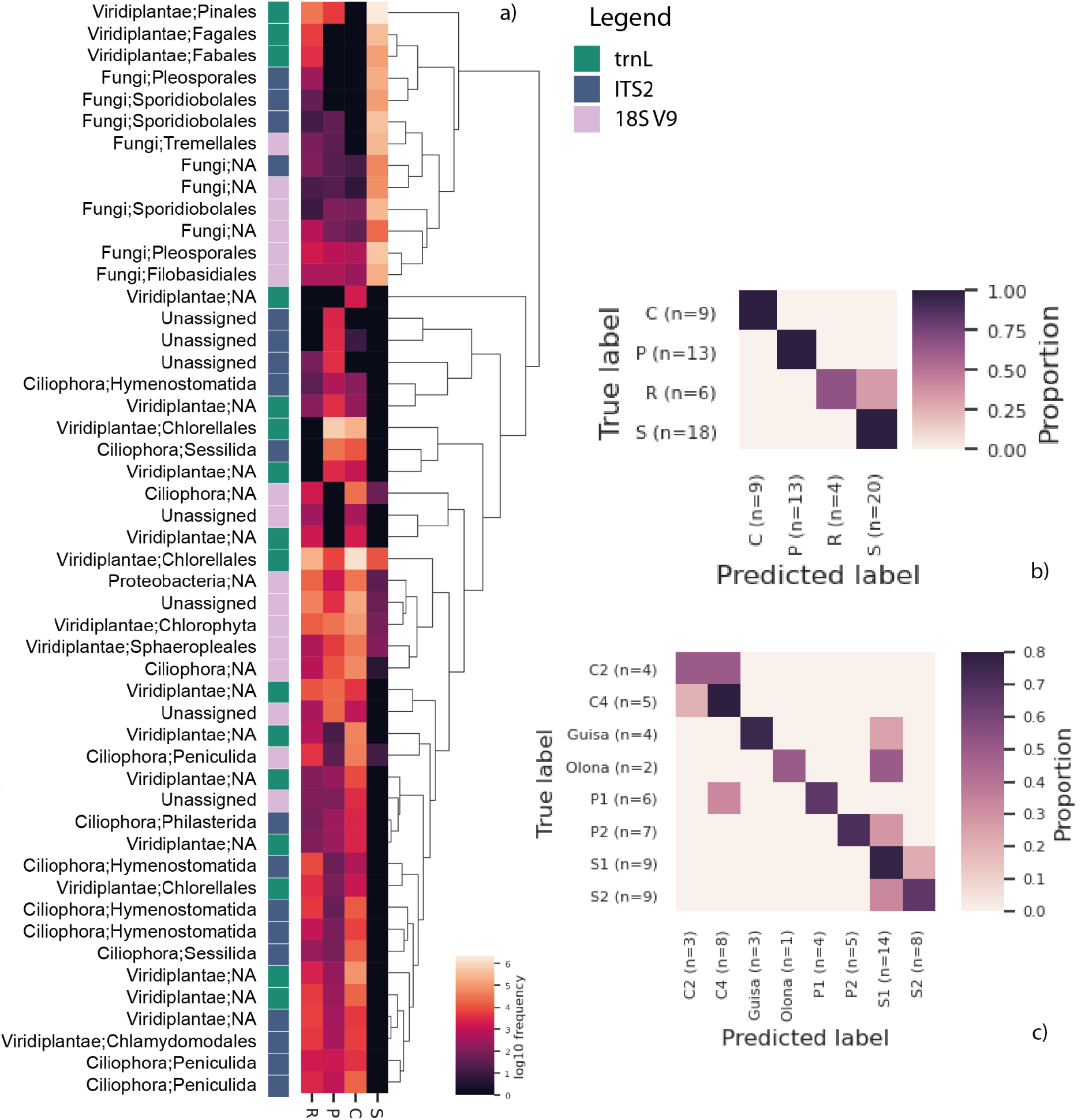
a) scatter plot representing ESVs with the additional information of marker and taxonomy (in particular: pink for V9 18S, green for trnL and blue for ITS2); scatter plot representing ESVs both using macro category sites (b) and sampling sites (c).

Considering V9 18S marker, the set of ESVs was able to discriminate between water sampling sites with high precision and recalling, difficulties remained in the classification of different air sites, but without bias in defining the macro category air site (S). Considering the use of data filtered, an optimal result was obtained considering Eukaryota sequences (Figure 6c), obtaining a higher recall. Using only ESVs (Figure 6b) or Metazoa sequences (Figure 6d), the macro category site prediction resulted in a lower performance, in particular regarding Rivers and Air sites prediction for sequences, and P and Air sites for Metazoa.

The trend described above was quite similar also for the other two marker regions. In detail, ITS2 sequences and Fungi worked well considering macro category prediction; predicting sampling sites using ESVs, instead, did not obtain optimal results, in particular for sites belonging to Guisa and Olona and P canals. Air sites, instead, were not correctly classified considering the division into the two sites, but the macro category was maintained (for details, see Supplementary S10).

Overall, machine learning analysis considering trnL markers reached good results, both considering ESVs and data filtered by taxa. Macro category prediction was reached. Streptophyta filtering showed difficulties in distinguishing River and Air sites. In general, the recalling was high (for details, see Supplementary S10).

Considering the information obtained by the three marker region sequencing, we decided to integrate all the data obtained from the ESVs calculated and run the machine learning classification considering all the ESVs obtained from the nine runs. The results are shown in Figure 6, representing the scatter plots obtained considering sampling sites and macro category prediction (Fig. 6b and c). In addition, the heatmap with the ESVs selected was represented, accompanied by the taxonomy at the Order rank (Fig. 6a). Also in this case, it is possible to observe a clear distinction between water and air medium type, a clear discrimination of canal water sites (C and P).

## Discussion

In this study we have tested the reliability of DNA metabarcoding in capturing the environmental effects on eukaryotic diversity of a mega-event. Based on the data obtained from the study, we tried to answer the following questions: is it possible to use DNA metabarcoding to track biodiversity communities in a mega-event context? Which are the pros and cons of using multi-marker strategies, considering the absence of common procedures and the issues related to the taxonomy assignment? Lastly, can machine learning procedures help in predicting sample origin, overcoming the taxonomic gap?

According to Bayraktarov et al. 2019, a common trend in most eDNA studies is the accumulation of data following the widespread opinion that “the more data there is, the better” (van Dorst et al., 2014). Nevertheless, this statement may be true only in specific contexts, since it disproportionately prioritizes data quantity over data quality. The value of data collected depends on how effective they are in achieving the solution to the key problem addressed (e.g., improving environmental management, Field et al., 2005, Miller et al., 2019).

Currently, DNA metabarcoding is the only reliable solution to collect large scale biodiversity data, as stated in the review paper of Cordier et al. 2021. The first question is if it is possible to use it as a routine tool for environmental biomonitoring. Several scientific studies demonstrated the feasibility of applying DNA metabarcoding in monitoring strategies, implementing both wet lab and bioinformatic pipelines in their workflows (Zafeiropoulos et al., 2020).

In this study, the event EXPO2015 provided an ideal system to test the effectiveness of DNA metabarcoding as a biomonitoring tool to check biodiversity variations in a critical ecosystem. The area is located in a highly urbanized environment, close to large suburban parks. These areas are proved to be an important reservoir of biodiversity, especially larger parks that could contribute more to the conservation of biodiversity (Cornelis and Hermy, 2004; McKinney, 2008; Beninde et al. 2015). Preserving these habitats is a fundamental point in the conservation of biodiversity, especially in a fragile context such as the urban one. For these reasons, validating eDNA metabarcoding tools is pivotal to monitor environments exposed to changes that could burden their equilibrium.

Going back to our first question, the capacity of our tool to detect variations is really powerful. Our results clearly showed a difference among the different sites (Figure 2-3-4, section “c” and “d”), but not a strong difference among the most distant sampling dates, considering both types of medium (Supplementary S6).

At the same time, the data collected by air and water were very different, both considering ESVs and taxa detected, as we can state by statistical analysis performed for all the markers investigated (see Supplementary S5 for details). Further, it was possible to individuate a fingerprint that made P sites different from C sites and rivers sites. In particular, PCoA analysis (Figure 2-3-4, section “d”) showed a clusterization among sites belonging to these three categories, identifying patterns of biodiversity that characterized distinct regions of the EXPO area, supported also by statistical tests and obtained considering both ESVs and taxonomic assignment (Figure 2-3-4, section “d”).

The approach addressed above opens our discussion to another question: is DNA metabarcoding a valid taxonomic identification tool? Several research papers demonstrated its application in diet characterization, water assessments, pollen identification and many other relevant fields (Porter et al., 2018; Deiner et al., 2017; Zhang et al., 2020). The taxonomic assignment step is still a delicate phase, as there are still no well-defined standards for each marker used in metabarcoding studies (Porter et al., 2018). In particular, the choice of the marker is still a compromise between two main aspects: i) the length of the DNA region that can be sequenced and, as a consequence, the genetic information that can be obtained; ii) the reference databases completeness and accuracy.

The first mostly regards the fact that any kind of matrix has its own characteristic. The key aspect to keep in mind is, usually, the DNA degradation, that can affect the reliability of the study. Shorter fragments are more likely to be detected, considering for example diets characterization or water assessments, where DNA is exposed to multiple degrading sources, like chemical compounds (Deiner et al., 2017) and temperature (Krehenwinkel et al., 2018). At the same time, the marker chosen will influence taxa detection (Deiner et al., 2017): a gap of references recorded versus the known biodiversity exists for several relevant taxa, impeding a complete and correct taxonomic assignment (Cordier et al., 2021; Weigand et al., 2019; McGee et al., 2019). If this aspect is not considered, important biases could be included into experiments, leading to misinterpretations and excluding crucial information.

The markers that we evaluated as suitable for our experiments are a compromise among all these issues. Selecting short length markers (of about 150-200 base pairs), such as 18S V9 and the intron region of trnL, allowed us to collect a great number of information about eukaryotic and plant groups, even considering the highly degraded matrices we collected in our sampling campaign. Similarly, the longer region of the internal transcribed spacer ITS2 represented a good trade-off between low length variation and universality of primer sites (Nilsson et al., 2018), thus providing an overview of the Fungi Kingdom. In order to completely explore the potential of DNA metabarcoding, we decided to show not only the taxonomic assignments of ESVs, but also to include the analyses evaluating the ability of each marker to reach the taxa group for which they are recommended (Porter et al., 2018; Deiner et al., 2017), considering also researches already conducted (e.g. for trnL Quemere et al., 2013; 18S V9 Fernández-Álvarez et al. 2018; ITS2 Banchi et al., 2018). Sunburst plots of taxonomic distribution of taxa detected are shown in Figure 2-3-4 section “b”. The category of Metazoa, Plants and Fungi was extracted to show the balance of variations across sites in Figure 2-3-4 section “c”.

Some critical issues that emerged in our study are still at the center of the current scientific debate. Despite the huge amount of data obtained from the sampling campaign, taxonomy assignment remains a difficult task. In general, the most of V9 18S sequences were assigned to unicellular eukaryotes taxa, followed by Fungi, Viridiplantae and Metazoa, with a 20% of Unassigned sequences. Considering ITS2 sequences, the majority were assigned to Fungi, followed by Unicellular Eukaryotes taxa, with 11% of Unassigned sequences. Lastly, plastid trnL intron sequences resulted in Streptophyta assignments, with no Unassigned sequences. Overall, exploring taxonomy results helped us to consider sequencing outputs from different points of view. In particular, interactive sunburst in Supplementary S7 were created to enable a correct comprehension of the data obtained. Basically, there are issues with the standardization of taxonomy through different taxonomic groups (in particular Unicellular Eukaryotes and Viridiplantae). Knowing the difficulties of markers to reach genus or species ranks, it happened to investigate diversity considering for example families or orders gaps into the description of taxonomy that will not allow the data to be interpreted correctly.

For this reason, we decided to evaluate both biodiversity and machine learning analysis considering not only the taxa of interest, but also the genetic information that we obtained. In general, the analysis was coherent considering all the markers used, also subgrouping the ESVs based on particular taxa. But for the sake of interpretability and standardization, we believe that a focus on ESVs without the taxonomic assignment must be taken into account for a reliable and correct analysis of DNA metabarcoding data.

Overall, PCoA (Figure 2-3-4 section “d”) clearly showed a significant structuration based on sampling site, with Internal sites (P1-P2) cluster closely as External sites (C1-C2) and Rivers (OL-GU), demonstrating that ESVs composition could be the key to identify site types among the EXPO area (rivers, sites C and sites P), overcoming the gap in reference databases.

Aside from the spatial information, we explored the effect of the sampling month (see Supplementary S6 for details). We think that the effect of Site was predominant, though samples belonging to the same site and period of sampling were much more similar, suggesting the presence of a fingerprint due to both spatial and period. As the difficulties related to collecting samples during the EXPO event, we cannot ascertain the importance of sampling time. But, considering several previous works related to temporal biomonitoring with DNA metabarcoding, we do not exclude time effects (Deiner et al., 2017; Pawlowski et al., 2018; Ruppert et al., 2019; Porter et al., 2018).

In general, our strategy suggests that the molecular information collected during the sample campaign was universally different in the sampling area and this trend was observed for all the three genetic surveys that we performed. The additional analysis carried out considering only the taxonomy demonstrated the strength of information collected.

Our last evaluation took into account if the DNA metabarcoding can be applied for predictive purpose. For this task, we used a machine learning approach both considering ESVs and taxa assigned.

We confirmed the biodiversity analysis conducted for all the markers. In addition, results indicated the use of sequences can be predictive, passing the taxonomy assignment that can be misleading. At the same time, a filter based on specific kingdoms suggests a peculiar structure for each taxa explored. Regarding this point, we are perfectly aware of the complexity of the communities that we analysed, but the recall is high, considering the vicinity of sampling sites and the medium of investigation.

In general, results obtained from machine learning classification showed three main aspects: the importance of sequences as a baseline pattern information of sites, the strength of the patterns considering also different taxonomic levels of analysis and, lastly, the optimization of the classification considering the macro site category. The use of taxonomy filtering for machine learning demonstrated the role of molecular fingerprinting, suggesting that the method can also be applied without reaching specific taxa information. This fact suggests two things: the first one is that DNA metabarcoding with middle-short region can be used for finding molecular fingerprinting in large-analysis and, in some cases, also taxonomy fingerprinting can be obtained and exploit (unfortunately, as we mentioned above, this really depends on the molecular marker used and the reference databases used for the assignment) (Schloss and Westcott, 2011). The second, instead, demonstrated the difficulties in using DNA metabarcoding in reaching species-level information.

We think that the complexity should not be underestimated. Considering the data that we showed and the results related, the level of investigation may be very different, allowing researchers to answer several questions. The type of matrices, sampling method and marker used may lead to a real selection of communities under studies.

From the end of EXPO2015, no alien species were detected by state control agencies (ARPA) next to the exposition area. From the biomonitoring point of view, it is clear that advances in collecting data and contributing to public repositories could make a difference in interpreting these results. However, large amounts of biodiversity data may be useful for the generation of hypotheses (Bayraktarov et al., 2019). In the last few years, an increase of publications of ‘DNA metabarcoding’ in monitoring, biosurveillance and species invasions was observed (Piper et al., 2019). Though the advent of new sequencing technologies bring us the possibility to collect longer reads, therefore more genetic information, DNA metabarcoding with mid-short marker genes is still an important methodology in biodiversity assessment (Piper et al., 2019). From the biomonitoring point view, traces of ESVs could be informative alone, in order to study patterns and without focusing on specific taxonomic groups, for which it is possible to implement taxa-specific markers (Elbrecht et al., 2019) or classic monitoring strategies (Cordier et al., 2021).

DNA metabarcoding is nowadays widely used for very different purposes. Several research papers demonstrated its applicability into the monitoring of biodiversity. In this research paper, we wanted to highlight not only the power, but also the limitations that have to be considered in order to manage the data and to give a conscious interpretation of data generated. As a genomic approach, limitations can be due to both markers chosen and the molecular information registered into the reference databases. Despite these issues, we demonstrated that the power of DNA metabarcoding is related not only to the molecular fingerprint obtained with sequencing ESVs, but also to the ability to collect a large amount of data, achieving a sort of freeze frame of the environment under study. For these reasons, bioinformatics and post-processing analysis is still a pivotal process. Mining information from genomic data is still an important task, not without difficulties, and in this context collecting information and submitting datasets to reference databases will only ameliorate the comprehension of biodiversity all around the world, implementing both our current knowledge and future research. Considering the trends related to open science and our ability to sequence and produce data, data mining approaches (e.g. machine learning) will become more and more important, helping in disentangling high amounts of data, detecting biodiversity patterns and integrating additional information that give an edge to future studies.

## Supporting information

Supplementary files directory

## Acknowledgements

We want to thank all the students who have worked to collect and analyse part of this data over the years. Without their work it would not have been possible to have such a large amount of data available. We also want to thank the company Expo 2015 S.p.A for showing its willingness to collaborate in this study by allowing continuous sampling.

Partial funding for the data collection and management is provided in the framework of LifeWatch, which is a landmark European Research Infrastructure within the European Strategy Forum on Research (ESFRI) roadmap.

## Data Accessibility

The dataset generated for this study was submitted to the EBI metagenomics portal (https://www.ebi.ac.uk/metagenomics/; Study ID: PRJEB45249) and it will be freely available upon paper submission and acceptance.

## Author Contributions

Giulia Agostinetto, Antonia Bruno and Anna Sandionigi analyzed the data and wrote the paper. Alberto Brusati, Caterina Manzari, Alice Chiodi and Eleonora Siani contributed to the experimental procedures and analytical tools. Maurizio Casiraghi conceived the idea. All authors listed have made a substantial, direct and intellectual contribution to the work, and approved it for publication.

## Footnotes

1 Expo 2015 S.p.A (2018) Expo Milano 2015 Official Report (http://www.expo2015.org/wp-content/uploads/2019/10/OFFICIAL%20REPORT%20EXPO%20MILANO%202015-PDF-ENG.pdf)

## References

Agostinetto, G., Brusati, A., Sandionigi, A., Chahed Adam, Parladori Elena, Bachir, B., Antonia, B., Dario, P., & Maurizio, C. (2021). Supporting data for “ExTaxsI: An exploration tool of biodiversity molecular data” (p. 1 GB) [Data set]. GigaScience Database.s

Agostinetto, G., Sandionigi, A., Chahed, A., Brusati, A., Parladori, E., Balech, B., Bruno, A., Pescini, D., & Casiraghi, M. (2020). ExTaxsI: an exploration tool of biodiversity molecular data. BioRxiv.

Alberdi, A., Aizpurua, O., Gilbert, M. T. P., & Bohmann, K. (2018). Scrutinizing key steps for reliable metabarcoding of environmental samples. Methods in Ecology and Evolution, 9(1), 134–147.

Anderson, M. J. (2001). Permutation tests for univariate or multivariate analysis of variance and regression. Canadian Journal of Fisheries and Aquatic Sciences, 58(3), 626–639.

Baird, D. J., & Hajibabaei, M. (2012). Biomonitoring 2.0: A new paradigm in ecosystem assessment made possible by next-generation DNA sequencing: NEWS AND VIEWS: OPINION. Molecular Ecology, 21(8), 2039–2044.

Banchi, E., Ametrano, C. G., Stanković, D., Verardo, P., Moretti, O., Gabrielli, F., Lazzarin, S., Borney, M. F., Tassan, F., & Tretiach, M. (2018). DNA metabarcoding uncovers fungal diversity of mixed airborne samples in Italy. PloS One, 13(3), e0194489.

Bayraktarov, E., Ehmke, G., O’connor, J., Burns, E. L., Nguyen, H. A., McRae, L., Possingham, H. P., & Lindenmayer, D. B. (2019). Do big unstructured biodiversity data mean more knowledge? Frontiers in Ecology and Evolution, 6, 239.

Beninde, J., Veith, M., & Hochkirch, A. (2015). Biodiversity in cities needs space: A meta-analysis of factors determining intra-urban biodiversity variation. Ecology Letters, 18(6), 581–592.

Blaalid, R., Kumar, S., Nilsson, R. H., Abarenkov, K., Kirk, P. M., & Kauserud, H. (2013). ITS 1 versus ITS 2 as DNA metabarcodes for fungi. Molecular Ecology Resources, 13(2), 218–224.

Bokulich, N. A., Dillon, M. R., Bolyen, E., Kaehler, B. D., Huttley, G. A., & Caporaso, J. G. (2018). q2-sample-classifier: Machine-learning tools for microbiome classification and regression. Journal of Open Research Software, 3(30).

Bolyen, E., Rideout, J. R., Dillon, M. R., Bokulich, N. A., Abnet, C. C., Al-Ghalith, G. A., Alexander, H., Alm, E. J., Arumugam, M., & Asnicar, F. (2019). Reproducible, interactive, scalable and extensible microbiome data science using QIIME 2. Nature Biotechnology, 37(8), 852–857.

Bonada, N., Prat, N., Resh, V. H., & Statzner, B. (2006). Developments in aquatic insect biomonitoring: A comparative analysis of recent approaches. Annu. Rev. Entomol., 51, 495–523.

Bongaarts, J. (2019). IPBES, 2019. Summary for policymakers of the global assessment report on biodiversity and ecosystem services of the Intergovernmental Science-Policy Platform on Biodiversity and Ecosystem Services. Population and Development Review, 45(3), 680–681.

Boyer, F., Mercier, C., Bonin, A., Le Bras, Y., Taberlet, P., & Coissac, E. (2016). obitools: A unix-inspired software package for DNA metabarcoding. Molecular Ecology Resources, 16(1), 176–182.

Bush, A., Compson, Z. G., Monk, W. A., Porter, T. M., Steeves, R., Emilson, E., Gagne, N., Hajibabaei, M., Roy, M., & Baird, D. J. (2019). Studying ecosystems with DNA metabarcoding: Lessons from biomonitoring of aquatic macroinvertebrates. Frontiers in Ecology and Evolution, 7, 434.

Callahan, B. J., McMurdie, P. J., & Holmes, S. P. (2017). Exact sequence variants should replace operational taxonomic units in marker-gene data analysis. The ISME Journal, 11(12), 2639–2643.

Callahan, B. J., McMurdie, P. J., Rosen, M. J., Han, A. W., Johnson, A. J. A., & Holmes, S. P. (2016). DADA2: High-resolution sample inference from Illumina amplicon data. Nature Methods, 13(7), 581–583.

Capo, E., Spong, G., Königsson, H., & Byström, P. (2020). Effects of filtration methods and water volume on the quantification of brown trout (Salmo trutta) and Arctic char (Salvelinus alpinus) eDNA concentrations via droplet digital PCR. Environmental DNA, 2(2), 152–160.

Chariton, A. A., Stephenson, S., Morgan, M. J., Steven, A. D., Colloff, M. J., Court, L. N., & Hardy, C. M. (2015). Metabarcoding of benthic eukaryote communities predicts the ecological condition of estuaries. Environmental Pollution, 203, 165–174.

Cheng, S., Melkonian, M., Smith, S. A., Brockington, S., Archibald, J. M., Delaux, P.-M., Li, F.-W., Melkonian, B., Mavrodiev, E. V., & Sun, W. (2018). 10KP: A phylodiverse genome sequencing plan. Gigascience, 7(3), giy013.

Comtet, T., Sandionigi, A., Viard, F., & Casiraghi, M. (2015). DNA (meta) barcoding of biological invasions: A powerful tool to elucidate invasion processes and help managing aliens. Biological Invasions, 17(3), 905–922.

Cordier, T., Alonso-Sáez, L., Apothéloz-Perret-Gentil, L., Aylagas, E., Bohan, D. A., Bouchez, A., Chariton, A., Creer, S., Frühe, L., Keck, F., Keeley, N., Laroche, O., Leese, F., Pochon, X., Stoeck, T., Pawlowski, J., & Lanzén, A. (2021). Ecosystems monitoring powered by environmental genomics: A review of current strategies with an implementation roadmap. Molecular Ecology, 30(13), 2937–2958.

Cornelis, J., & Hermy, M. (2004). Biodiversity relationships in urban and suburban parks in Flanders. Landscape and Urban Planning, 69(4), 385–401.

Cowart, D. A., Pinheiro, M., Mouchel, O., Maguer, M., Grall, J., Miné, J., & Arnaud-Haond, S. (2015). Metabarcoding is powerful yet still blind: A comparative analysis of morphological and molecular surveys of seagrass communities. PloS One, 10(2), e0117562.

Cristescu, M. E., & Hebert, P. D. (2018). Uses and misuses of environmental DNA in biodiversity science and conservation. Annual Review of Ecology, Evolution, and Systematics, 49, 209–230.

Curry, C. J., Gibson, J. F., Shokralla, S., Hajibabaei, M., & Baird, D. J. (2018). Identifying North American freshwater invertebrates using DNA barcodes: Are existing COI sequence libraries fit for purpose? Freshwater Science, 37(1), 178–189.

Deiner, K., Bik, H. M., Mächler, E., Seymour, M., Lacoursière-Roussel, A., Altermatt, F., Creer, S., Bista, I., Lodge, D. M., & De Vere, N. (2017). Environmental DNA metabarcoding: Transforming how we survey animal and plant communities. Molecular Ecology, 26(21), 5872–5895.

Dequiedt, S., Saby, N. P. A., Lelievre, M., Jolivet, C., Thioulouse, J., Toutain, B., Arrouays, D., Bispo, A., Lemanceau, P., & Ranjard, L. (2011). Biogeographical patterns of soil molecular microbial biomass as influenced by soil characteristics and management. Global Ecology and Biogeography, 20(4), 641–652.

Edgcomb, V., Orsi, W., Bunge, J., Jeon, S., Christen, R., Leslin, C., Holder, M., Taylor, G. T., Suarez, P., & Varela, R. (2011). Protistan microbial observatory in the Cariaco Basin, Caribbean. I. Pyrosequencing vs Sanger insights into species richness. The ISME Journal, 5(8), 1344–1356.

Elbrecht, V., Braukmann, T. W., Ivanova, N. V., Prosser, S. W., Hajibabaei, M., Wright, M., Zakharov, E. V., Hebert, P. D., & Steinke, D. (2019). Validation of COI metabarcoding primers for terrestrial arthropods. PeerJ, 7, e7745.

Fahner, N. A., Shokralla, S., Baird, D. J., & Hajibabaei, M. (2016). Large-scale monitoring of plants through environmental DNA metabarcoding of soil: Recovery, resolution, and annotation of four DNA markers. PloS One, 11(6), e0157505.

Fernández-Álvarez, F. Á., Machordom, A., García-Jiménez, R., Salinas-Zavala, C. A., & Villanueva, R. (2018). Predatory flying squids are detritivores during their early planktonic life. Scientific Reports, 8(1), 1–12.

Field, S. A., Tyre, A. J., & Possingham, H. P. (2005). Optimizing allocation of monitoring effort under economic and observational constraints. The Journal of Wildlife Management, 69(2), 473–482.

Frontalini, F., Greco, M., Di Bella, L., Lejzerowicz, F., Reo, E., Caruso, A., Cosentino, C., Maccotta, A., Scopelliti, G., & Nardelli, M. P. (2018). Assessing the effect of mercury pollution on cultured benthic foraminifera community using morphological and eDNA metabarcoding approaches. Marine Pollution Bulletin, 129(2), 512–524.

Harrison, J. P., Chronopoulou, M., Salonen, I., Jilbert, T., & Koho, K. (2021). 16S and 18S rRNA gene metabarcoding provide congruent information on the responses of sediment communities to eutrophication. Frontiers in Marine Science, 8, 862.

Herbold, C. W., Pelikan, C., Kuzyk, O., Hausmann, B., Angel, R., Berry, D., & Loy, A. (2015). A flexible and economical barcoding approach for highly multiplexed amplicon sequencing of diverse target genes. Frontiers in Microbiology, 6.

Hirst, J. (1952). An automatic volumetric spore trap. Annals of Applied Biology, 39(2), 257–265.

Jamwal, P. S., Bruno, A., Galimberti, A., Magnani, D., Krupa, H., Casiraghi, M., & Loy, A. (2021). First assessment of eDNA-based detection approach to monitor the presence of Eurasian otter in southern Italy. Hystrix, the Italian Journal of Mammalogy.

Kanz, C., Aldebert, P., Althorpe, N., Baker, W., Baldwin, A., Bates, K., Browne, P., van den Broek, A., Castro, M., & Cochrane, G. (2005). The EMBL nucleotide sequence database. Nucleic Acids Research, 33(suppl_1), D29–D33.

Knights, D., Kuczynski, J., Koren, O., Ley, R. E., Field, D., Knight, R., DeSantis, T. Z., & Kelley, S. T. (2011). Supervised classification of microbiota mitigates mislabeling errors. The ISME Journal, 5(4), 570–573.

Krehenwinkel, H., Fong, M., Kennedy, S., Huang, E. G., Noriyuki, S., Cayetano, L., & Gillespie, R. (2018). The effect of DNA degradation bias in passive sampling devices on metabarcoding studies of arthropod communities and their associated microbiota. PLoS One, 13(1), e0189188.

Kruskal, W. H., & Wallis, W. A. (1952). Use of ranks in one-criterion variance analysis. Journal of the American Statistical Association, 47(260), 583–621.

Lallias, D., Hiddink, J. G., Fonseca, V. G., Gaspar, J. M., Sung, W., Neill, S. P., Barnes, N., Ferrero, T., Hall, N., & Lambshead, P. J. D. (2015). Environmental metabarcoding reveals heterogeneous drivers of microbial eukaryote diversity in contrasting estuarine ecosystems. The ISME Journal, 9(5), 1208–1221.

Lanzén, A., Lekang, K., Jonassen, I., Thompson, E. M., & Troedsson, C. (2016). High-throughput metabarcoding of eukaryotic diversity for environmental monitoring of offshore oil-drilling activities. Molecular Ecology, 25(17), 4392–4406.

Magurran, A. E., Baillie, S. R., Buckland, S. T., Dick, J. M., Elston, D. A., Scott, E. M., Smith, R. I., Somerfield, P. J., & Watt, A. D. (2010). Long-term datasets in biodiversity research and monitoring: Assessing change in ecological communities through time. Trends in Ecology & Evolution, 25(10), 574–582.

Makiola, A., Compson, Z. G., Baird, D. J., Barnes, M. A., Boerlijst, S. P., Bouchez, A., Brennan, G., Bush, A., Canard, E., Cordier, T., Creer, S., Curry, R. A., David, P., Dumbrell, A. J., Gravel, D., Hajibabaei, M., Hayden, B., van der Hoorn, B., Jarne, P., … Bohan, D. A. (2020). Key Questions for Next-Generation Biomonitoring. Frontiers in Environmental Science, 7, 197.

McGee, K. M., Robinson, C. V., & Hajibabaei, M. (2019). Gaps in DNA-Based Biomonitoring Across the Globe. Frontiers in Ecology and Evolution, 7, 337.

McKinney, M. L. (2008). Effects of urbanization on species richness: A review of plants and animals. Urban Ecosystems, 11(2), 161–176.

Miller, D. A., Pacifici, K., Sanderlin, J. S., & Reich, B. J. (2019). The recent past and promising future for data integration methods to estimate species’ distributions. Methods in Ecology and Evolution, 10(1), 22–37.

Nilsson, R. H., Anslan, S., Bahram, M., Wurzbacher, C., Baldrian, P., & Tedersoo, L. (2019). Mycobiome diversity: High-throughput sequencing and identification of fungi. Nature Reviews Microbiology, 17(2), 95–109.

Núñez, A., Amo de Paz, G., Ferencova, Z., Rastrojo, A., Guantes, R., García, A. M., Alcamí, A., Gutiérrez-Bustillo, A. M., & Moreno, D. A. (2017). Validation of the hirst-type spore trap for simultaneous monitoring of prokaryotic and eukaryotic biodiversities in urban air samples by next-generation sequencing. Applied and Environmental Microbiology, 83(13), e00472–17.

Pawlowski, J., Kelly-Quinn, M., Altermatt, F., Apothéloz-Perret-Gentil, L., Beja, P., Boggero, A., Borja, A., Bouchez, A., Cordier, T., & Domaizon, I. (2018). The future of biotic indices in the ecogenomic era: Integrating (e) DNA metabarcoding in biological assessment of aquatic ecosystems. Science of the Total Environment, 637, 1295–1310.

Pimm, S. L., Alibhai, S., Bergl, R., Dehgan, A., Giri, C., Jewell, Z., Joppa, L., Kays, R., & Loarie, S. (2015). Emerging Technologies to Conserve Biodiversity. Trends in Ecology & Evolution, 30(11), 685–696.

Piper, A. M., Batovska, J., Cogan, N. O., Weiss, J., Cunningham, J. P., Rodoni, B. C., & Blacket, M. J. (2019). Prospects and challenges of implementing DNA metabarcoding for high-throughput insect surveillance. GigaScience, 8(8), giz092.

Porter, T. M., & Hajibabaei, M. (2018). Scaling up: A guide to high-throughput genomic approaches for biodiversity analysis. Molecular Ecology, 27(2), 313–338.

Quéméré, E., Hibert, F., Miquel, C., Lhuillier, E., Rasolondraibe, E., Champeau, J., Rabarivola, C., Nusbaumer, L., Chatelain, C., & Gautier, L. (2013). A DNA metabarcoding study of a primate dietary diversity and plasticity across its entire fragmented range. PloS One, 8(3), e58971.

Reavie, E. D., Jicha, T. M., Angradi, T. R., Bolgrien, D. W., & Hill, B. H. (2010). Algal assemblages for large river monitoring: Comparison among biovolume, absolute and relative abundance metrics. Ecological Indicators, 10(2), 167–177.

Rhie, A., McCarthy, S. A., Fedrigo, O., Damas, J., Formenti, G., Koren, S., Uliano-Silva, M., Chow, W., Fungtammasan, A., & Kim, J. (2021). Towards complete and error-free genome assemblies of all vertebrate species. Nature, 592(7856), 737–746.

Ruppert, K. M., Kline, R. J., & Rahman, M. S. (2019). Past, present, and future perspectives of environmental DNA (eDNA) metabarcoding: A systematic review in methods, monitoring, and applications of global eDNA. Global Ecology and Conservation, 17, e00547.

Schloss, P. D., & Westcott, S. L. (2011). Assessing and improving methods used in operational taxonomic unit-based approaches for 16S rRNA gene sequence analysis. Applied and Environmental Microbiology, 77(10), 3219–3226.

Shokralla, S., Spall, J. L., Gibson, J. F., & Hajibabaei, M. (2012). Next-generation sequencing technologies for environmental DNA research. Molecular Ecology, 21(8), 1794–1805.

Taberlet, P., Gielly, L., Pautou, G., & Bouvet, J. (1991). Universal primers for amplification of three non-coding regions of chloroplast DNA. Plant Molecular Biology, 17(5), 1105–1109.

Tauber, H. (1974). A static non-overload pollen collector. New Phytologist, 73(2), 359–369.

Taylor, H. R., & Gemmell, N. J. (2016). Emerging Technologies to Conserve Biodiversity: Further Opportunities via Genomics. Response to Pimm et al. Trends in Ecology & Evolution, 31(3), 171–172.

Thomsen, P. F., & Willerslev, E. (2015). Environmental DNA–An emerging tool in conservation for monitoring past and present biodiversity. Biological Conservation, 183, 4–18.

Toju, H., Tanabe, A. S., Yamamoto, S., & Sato, H. (2012). High-coverage ITS primers for the DNA-based identification of ascomycetes and basidiomycetes in environmental samples. PloS One, 7(7), e40863.

Tommasi, N., Biella, P., Guzzetti, L., Lasway, J. V., Njovu, H. K., Tapparo, A., Agostinetto, G., Peters, M. K., Steffan-Dewenter, I., & Labra, M. (2021). Impact of land use intensification and local features on plants and pollinators in Sub-Saharan smallholder farms. Agriculture, Ecosystems & Environment, 319, 107560.

Trebitz, A. S., Hoffman, J. C., Darling, J. A., Pilgrim, E. M., Kelly, J. R., Brown, E. A., Chadderton, W. L., Egan, S. P., Grey, E. K., & Hashsham, S. A. (2017). Early detection monitoring for aquatic non-indigenous species: Optimizing surveillance, incorporating advanced technologies, and identifying research needs. Journal of Environmental Management, 202, 299–310.

Valsecchi, E., Coppola, E., Pires, R., Parmegiani, A., Casiraghi, M., Galli, P., & Bruno, A. (2021). Newly developed ad hoc molecular assays shows how eDNA can witness and anticipate the monk seal recolonization of central Mediterranean. BioRxiv.

van der Heyde, M., Bunce, M., Wardell-Johnson, G., Fernandes, K., White, N. E., & Nevill, P. (2020). Testing multiple substrates for terrestrial biodiversity monitoring using environmental DNA metabarcoding. Molecular Ecology Resources, 20(3), 732–745.

van Dorst, J., Bissett, A., Palmer, A. S., Brown, M., Snape, I., Stark, J. S., Raymond, B., McKinlay, J., Ji, M., & Winsley, T. (2014). Community fingerprinting in a sequencing world. FEMS Microbiology Ecology, 89(2), 316–330.

Vasselon, V., Rimet, F., Tapolczai, K., & Bouchez, A. (2017). Assessing ecological status with diatoms DNA metabarcoding: Scaling-up on a WFD monitoring network (Mayotte island, France). Ecological Indicators, 82, 1–12.

Waterhouse, R. M., Adam-Blondon, A.-F., Agosti, D., Baldrian, P., Balech, B., Corre, E., Davey, R. P., Lantz, H., Pesole, G., & Quast, C. (2021). Recommendations for connecting molecular sequence and biodiversity research infrastructures through ELIXIR. F1000Research, 10(1238), 1238.

Weigand, H., Beermann, A. J.,Čiampor, F., Costa, F. O., Csabai, Z., Duarte, S., Geiger, M. F., Grabowski, M., Rimet, F., & Rulik, B. (2019a). DNA barcode reference libraries for the monitoring of aquatic biota in Europe: Gap-analysis and recommendations for future work. Science of the Total Environment, 678, 499–524.

Weigand, H., Beermann, A. J., Čiampor, F., Costa, F. O., Csabai, Z., Duarte, S., Geiger, M. F., Grabowski, M., Rimet, F., & Rulik, B. (2019b). DNA barcode reference libraries for the monitoring of aquatic biota in Europe: Gap-analysis and recommendations for future work. Science of the Total Environment, 678, 499–524.

Westfall, K. M., Therriault, T. W., & Abbott, C. L. (2020). A new approach to molecular biosurveillance of invasive species using DNA metabarcoding. Global Change Biology, 26(2), 1012–1022.

White, T. J., Bruns, T., Lee, S., & Taylor, J. (1990). Amplification and direct sequencing of fungal ribosomal RNA genes for phylogenetics. PCR Protocols: A Guide to Methods and Applications, 18(1), 315–322.

Zafeiropoulos, H., Viet, H. Q., Vasileiadou, K., Potirakis, A., Arvanitidis, C., Topalis, P., Pavloudi, C., & Pafilis, E. (2020). PEMA: a flexible Pipeline for Environmental DNA Metabarcoding Analysis of the 16S/18S ribosomal RNA, ITS, and COI marker genes. GigaScience, 9(3), giaa022.

Zhang, S., Zhao, J., & Yao, M. (2020). A comprehensive and comparative evaluation of primers for metabarcoding eDNA from fish. Methods in Ecology and Evolution, 11(12), 1609–1625.

Zimmermann, J., Glöckner, G., Jahn, R., Enke, N., & Gemeinholzer, B. (2015). Metabarcoding vs. Morphological identification to assess diatom diversity in environmental studies. Molecular Ecology Resources, 15(3), 526–542.

