## Supplementary data for "Dealing with the promise of metabarcoding in mega-event biomonitoring: EXPO2015 unedited data"

### S1. Metadata files

Files included information regarding samples, in particular type of medium, sampling site and macro category site.
Files (TSV format):

- S1a: metadata v9 18S;
- S1b: metadata ITS2;
- S1c: metadata trnL

### S2. qPCR quantification of target DNA

Quantitative real-time PCR (qPCR) assays were performed with AB 7500 (Applied Biosystem). qPCR conditions included an initial denaturation at 95°C for 10 min, followed by 40 cycles of denaturation at 95°C for 15 s and annealing-elongation for 1 min at 55°C. Real-Time PCR was set up with 2X SsoFast EvaGreen Supermix with Low ROX [Bio-Rad S.r.l., Segrate (MI), Italy] in which EvaGreen was used as a detecting dye; a 10 µl reaction consisted of 5.0 µl SsoFast EvaGreen Supermix with Low ROX, 0.1 µl each 10 µmol/L primer solution, 2 µl DNA sample, and 2.8 µl of Milli-Q water. Primer sequences, targets, annealing temperatures, and references are given in Table SX. Standard curves were generated using 10-fold serial dilutions of positive controls and qPCR amplification efficiencies (E) were based on the following Equation (1):

$$E=10(-1/slope)-1$$

and R2 values (linearity) were 0.99. All samples and standards were run in triplicate. Negative controls were also tested in triplicate too for each amplification.
All the assays were followed by a dissociation stage and melting curves were obtained. Amplification data were collected and analyzed with the SDS 7500 Real-Time PCR System Software (Applied Biosystems). We reported all the data derived from qPCR as DNA copies of the target amplified, rather than Ct, normalized for the efficiency, following the formula

$$DNAcopies=E^\wedge(Ct1-Ct)$$

where E is the efficiency of amplification and Ct1 is the number of qPCR cycles required to detect a single target molecule (Bruno et al., 2017).

### S3. Table of primers used for each marker gene

| Primer | Region | Target | Sequence Reference |
|---|---|---|---|
| 1391F | 18SV9 | GTACACACCGCCCGTC | Edgcomb et al., 2011[1] |
| EukB | 18SV9 | TGATCCTTCTGCAGGTTCACCTAC | Edgcomb et al., 2011 |
| trnL-C | P6 Loop trnL | CGAAATCGGTAGACGCTACG | Taberlet et al., 1991[2] |

| | | | |
|---|---|---|---|
| trnL-H | P6 Loop trnL | CCATTGAGTCTCTGCACCTATC | Taberlet et al., 1991 |
| ITS3KYO2 | ITS2 | GATGAAGAACGYAGYRAA | Toju et al., 2012[3] |
| ITS4 | ITS2 | TCCTCCGCTTATTGATATGC | White et al, 1990[4] |

[1]Edgcomb, Virginia, et al. "Protistan microbial observatory in the Cariaco Basin, Caribbean. I. Pyrosequencing vs Sanger insights into species richness." *The ISME journal* 5.8 (2011): 1344-1356.
[2]Taberlet, Pierre, et al. "Universal primers for amplification of three non-coding regions of chloroplast DNA." *Plant molecular biology* 17.5 (1991): 1105-1109.
[3]Toju, Hirokazu, et al. "High-coverage ITS primers for the DNA-based identification of ascomycetes and basidiomycetes in environmental samples." *PloS one* 7.7 (2012).
[4]White, Thomas J., et al. "Amplification and direct sequencing of fungal ribosomal RNA genes for phylogenetics." *PCR protocols: a guide to methods and applications* 18.1 (1990): 315-322.

#### S4. Percentage of ranks reached during the taxonomy assignment for each marker gene.

| Marker | Kingdom | Phylum | Class | Order | Family | Genus | Species |
|---|---|---|---|---|---|---|---|
| V9 (19304) | 23.68% | 33.61% | 24.40% | 26.23% | 24.08% | 19.18% | 12.98% |
| ITS2 (8471) | 72.73% | 78% | 71.76% | 66.31% | 61.54% | 51.63% | 36.90% |
| trnL (3630) | 100% | 57.82% | 11.26% | 22.36% | 21.51% | 14.52% | 4.84% |

#### S5a. Alpha diversity

#### V9 18S

| Group 1 | Group 2 | Sample size | H | P-value |
|---|---|---|---|---|
| Air | Water | 81 | 40.93 | < 0.001 |
| S1 | S2 | 34 | 0.17 | 0.68 |

| Group 1 | Group 2 | Sample size | H | P-value | Q-value |
|---|---|---|---|---|---|
| Water sites (C, P, R) | | 47 | 10.2 | 0.006 | / |
| Pairwise comparisons | | | | | |
| C | P | 37 | 10.3 | 0.001 | 0.004 |
| C | R | 26 | 2.02 | 0.15 | 0.23 |

| | | | | | |
|---|---|---|---|---|---|
| P | R | 31 | 1.4 | 0.23 | 0.23 |

| Group 1 | Group 2 | Sample size | H | P-value | Q-value |
|---|---|---|---|---|---|
| Water sites (C2, C4, GU, OL, P1, P2) | | 47 | 15.41 | **0.009** | / |
| Pairwise comparisons | | | | | |
| C2 | C4 | 16 | 1.23 | 0.26 | 0.44 |
| C2 | GU | 13 | 0 | 1 | 1 |
| C2 | OL | 11 | 0.32 | 0.57 | 0.71 |
| C2 | P1 | 17 | 0.53 | 0.46 | 0.7 |
| C2 | P2 | 18 | 5.3 | 0.02 | 0.07 |
| C4 | GU | 15 | 1.4 | 0.24 | 0.44 |
| C4 | OL | 13 | 4.66 | 0.03 | 0.09 |
| C4 | P1 | 19 | 6 | 0.01 | 0.07 |
| C4 | P2 | 20 | 9.94 | 0.001 | **0.02** |
| GU | OL | 10 | 0.4 | 0.52 | 0.71 |
| GU | P1 | 16 | 0.18 | 0.66 | 0.72 |
| GU | P2 | 17 | 2.58 | 0.1 | 0.27 |
| OL | P1 | 14 | 0.18 | 0.67 | 0.72 |
| OL | P2 | 15 | 2.06 | 0.15 | 0.32 |
| P1 | P2 | 21 | 5.73 | 0.01 | 0.07 |

### V9 18S - Eukaryota

| Group 1 | Group 2 | Sample size | H | P-value |
|---|---|---|---|---|
| Air | Water | 81 | 37.10 | **< 0.001** |
| S1 | S2 | 34 | 0.23 | 0.62 |

| Group 1 | Group 2 | Sample size | H | P-value | Q-value |
|---|---|---|---|---|---|
| Water sites (C, P, R) | | 47 | 11.87 | **0.002** | / |
| Pairwise comparisons | | | | | |
| C | P | 37 | 12 | 0.0005 | **0.001** |
| C | R | 26 | 2.18 | 0.14 | 0.19 |

| | | | | | |
|---|---|---|---|---|---|
| P | R | 31 | 1.72 | 0.19 | 0.19 |

| Group 1 | Group 2 | Sample size | H | P-value | Q-value |
|---|---|---|---|---|---|
| Water sites (C2, C4, GU, OL, P1, P2) | | 47 | 16.58 | **0.005** | / |
| Pairwise comparisons | | | | | |
| C2 | C4 | 16 | 1.12 | 0.29 | 0.48 |
| C2 | GU | 13 | 0.081 | 0.77 | 0.83 |
| C2 | OL | 11 | 0.32 | 0.57 | 0.71 |
| C2 | P1 | 17 | 0.95 | 0.33 | 0.49 |
| C2 | P2 | 18 | 7.14 | 0.007 | **0.05** |
| C4 | GU | 15 | 1.68 | 0.19 | 0.36 |
| C4 | OL | 13 | 3.42 | 0.06 | 0.19 |
| C4 | P1 | 19 | 6.4 | 0.01 | **0.05** |
| C4 | P2 | 20 | 10.42 | 0.001 | **0.01** |
| GU | OL | 10 | 0.41 | 0.52 | 0.71 |
| GU | P1 | 16 | 0.19 | 0.66 | 0.76 |
| GU | P2 | 17 | 2.27 | 0.13 | 0.28 |
| OL | P1 | 14 | 0.02 | 0.89 | 0.89 |
| OL | P2 | 15 | 2.88 | 0.09 | 0.22 |
| P1 | P2 | 21 | 6.07 | 0.01 | **0.05** |

### V9 18S - Metazoa

| Group 1 | Group 2 | Sample size | H | P-value |
|---|---|---|---|---|
| Air | Water | 81 | 31.36 | **< 0.001** |
| S1 | S2 | 34 | 0.95 | 0.33 |

| Group 1 | Group 2 | Sample size | H | P-value | Q-value |
|---|---|---|---|---|---|
| Water sites (C, P, R) | | 47 | 2.02 | 0.36 | / |
| Pairwise comparisons | | | | | |
| C | P | 37 | 1.36 | 0.24 | 0.4 |

| | | | | | |
|---|---|---|---|---|---|
| C | R | 26 | 0.16 | 0.69 | 0.7 |
| P | R | 31 | 1.24 | 0.26 | 0.4 |

| Group 1 | Group 2 | Sample size | H | P-value | Q-value |
|---|---|---|---|---|---|
| Water sites (C2, C4, GU, OL, P1, P2) | | 47 | 6.28 | 0.28 | / |
| Pairwise comparisons | | | | | |
| C2 | C4 | 16 | 4.49 | 0.03 | 0.41 |
| C2 | GU | 13 | 2.56 | 0.1 | 0.41 |
| C2 | OL | 11 | 0.32 | 0.57 | 0.91 |
| C2 | P1 | 17 | 0.03 | 0.84 | 0.91 |
| C2 | P2 | 18 | 0.1 | 0.75 | 0.91 |
| C4 | GU | 15 | 0.03 | 0.85 | 0.91 |
| C4 | OL | 13 | 0.21 | 0.64 | 0.91 |
| C4 | P1 | 19 | 2.67 | 0.1 | 0.41 |
| C4 | P2 | 20 | 3.19 | 0.7 | 0.41 |
| GU | OL | 10 | 0.12 | 0.7 | 0.91 |
| GU | P1 | 16 | 1.21 | 0.27 | 0.68 |
| GU | P2 | 17 | 1.98 | 0.16 | 0.48 |
| OL | P1 | 14 | 0.005 | 0.94 | 0.94 |
| OL | P2 | 15 | 0.61 | 0.43 | 0.91 |
| P1 | P2 | 21 | 0.04 | 0.83 | 0.91 |

### ITS2

| Group 1 | Group 2 | Sample size | H | P-value |
|---|---|---|---|---|
| Air | Water | 62 | 3.27 | 0.07 |
| S1 | S2 | 22 | 0.07 | 0.79 |

| Group 1 | Group 2 | Sample size | H | P-value | Q-value |
|---|---|---|---|---|---|
| Water sites (C, P, R) | | 43 | 11.96 | **0.002** | / |

| Pairwise comparisons | | | | | |
|---|---|---|---|---|---|
| C | P | 33 | 10.63 | 0.001 | **0.003** |
| C | R | 23 | 5.26 | 0.21 | **0.03** |
| P | R | 32 | 2.39 | 0.12 | 0.12 |

| Group 1 | Group 2 | Sample size | H | P-value | Q-value |
|---|---|---|---|---|---|
| Water sites (C2, C4, GU, OL, P1, P2) | | 43 | 13.01 | **0.02** | / |
| Pairwise comparisons | | | | | |
| C2 | C4 | 13 | 0.53 | 0.46 | 0.58 |
| C2 | GU | 11 | 3.33 | 0.07 | 0.17 |
| C2 | OL | 9 | 1.5 | 0.22 | 0.31 |
| C2 | P1 | 15 | 3.37 | 0.06 | 0.17 |
| C2 | P2 | 17 | 4.9 | 0.03 | 0.13 |
| C4 | GU | 14 | 3.7 | 0.05 | 0.17 |
| C4 | OL | 12 | 1.4 | 0.23 | 0.32 |
| C4 | P1 | 18 | 6.2 | 0.01 | 0.11 |
| C4 | P2 | 20 | 5.9 | 0.01 | 0.11 |
| GU | OL | 10 | 0.18 | 0.67 | 0.71 |
| GU | P1 | 16 | 2.64 | 0.1 | 0.22 |
| GU | P2 | 18 | 2.24 | 0.13 | 0.25 |
| OL | P1 | 14 | 0.32 | 0.57 | 0.66 |
| OL | P2 | 16 | 0.13 | 0.71 | 0.71 |
| P1 | P2 | 22 | 2.01 | 0.15 | 0.26 |

### ITS2 - Fungi

| Group 1 | Group 2 | Sample size | H | P-value |
|---|---|---|---|---|
| Air | Water | 35 | 3.15 | 0.076 |
| S1 | S2 | 22 | 0.04 | 0.84 |

| Group 1 | Group 2 | Sample size | H | P-value | Q-value |
|---|---|---|---|---|---|
| Water sites (C, P, R) | | 35 | 16.73 | **0.0002** | / |

| | | | | | |
|---|---|---|---|---|---|
| Pairwise comparisons | | | | | |
| C | P | 16 | 5.38 | 0.02 | **0.02** |
| C | R | 17 | 9.81 | 0.001 | **0.002** |
| P | R | 27 | 10.93 | 0.0009 | 0.002 |

| Group 1 | Group 2 | Sample size | H | P-value | Q-value |
|---|---|---|---|---|---|
| Water sites (C2, C4, GU, OL, P1, P2) | | 35 | 21.09 | **0.0007** | / |
| Pairwise comparisons | | | | | |
| C2 | C4 | 8 | 0.2 | 0.65 | 0.65 |
| C2 | GU | 10 | 6.86 | 0.009 | **0.04** |
| C2 | OL | 9 | 3.87 | 0.049 | 0.8 |
| C2 | P1 | 13 | 0.92 | 0.38 | 0.36 |
| C2 | P2 | 15 | 4.17 | 0.04 | 0.08 |
| C4 | GU | 8 | 5 | 0.02 | 0.06 |
| C4 | OL | 7 | 2.58 | 0.11 | 0.15 |
| C4 | P1 | 11 | 1.06 | 0.3 | 0.35 |
| C4 | P2 | 13 | 5.37 | 0.02 | 0.06 |
| GU | OL | 9 | 4.37 | 0.03 | 0.08 |
| GU | P1 | 13 | 8.6 | 0.003 | **0.02** |
| GU | P2 | 15 | 9.54 | 0.002 | **0.02** |
| OL | P1 | 12 | 3.2 | 0.07 | 0.11 |
| OL | P2 | 14 | 1.48 | 0.22 | 0.3 |
| P1 | P2 | 18 | 5.46 | 0.01 | 0.06 |

### trnL

| Group 1 | Group 2 | Sample size | H | P-value |
|---|---|---|---|---|
| Air | Water | 73 | 42.88 | **<0.001** |
| S1 | S2 | 30 | 0.36 | 0.55 |

| Group 1 | Group 2 | Sample size | H | P-value | Q-value |
|---|---|---|---|---|---|
| Water sites (C, P, R) | | 40 | 5.06 | 0.08 | / |

| Pairwise comparisons | | | | | |
|---|---|---|---|---|---|
| C | P | 30 | 3.64 | 0.05 | 0.08 |
| C | R | 25 | 3.66 | 0.05 | 0.08 |
| P | R | 25 | 0.04 | 0.84 | 0.08 |

| Group 1 | Group 2 | Sample size | H | P-value | Q-value |
|---|---|---|---|---|---|
| Water sites (C2, C4, GU, OL, P1, P2) | | 40 | 6 | 0.3 | / |
| Pairwise comparisons | | | | | |
| C2 | C4 | 15 | 0.35 | 0.55 | 0.76 |
| C2 | GU | 12 | 2.07 | 0.15 | 0.56 |
| C2 | OL | 10 | 0.4 | 0.52 | 0.76 |
| C2 | P1 | 12 | 0.92 | 0.33 | 0.72 |
| C2 | P2 | 15 | 2.34 | 0.12 | 0.56 |
| C4 | GU | 15 | 4.26 | 0.04 | 0.56 |
| C4 | OL | 13 | 0.59 | 0.44 | 0.76 |
| C4 | P1 | 15 | 1.4 | 0.24 | 0.71 |
| C4 | P2 | 18 | 2.25 | 0.13 | 0.56 |
| GU | OL | 10 | 1.13 | 0.28 | 0.71 |
| GU | P1 | 12 | 0.02 | 0.87 | 0.87 |
| GU | P2 | 15 | 0.05 | 0.81 | 0.87 |
| OL | P1 | 10 | 0.18 | 0.67 | 0.77 |
| OL | P2 | 13 | 0.48 | 0.48 | 0.76 |
| P1 | P2 | 15 | 0.22 | 0.64 | 0.77 |

### trnL - Streptophyta

| Group 1 | Group 2 | Sample size | H | P-value |
|---|---|---|---|---|
| Air | Water | 75 | 10.32 | **0.001** |
| S1 | S2 | 30 | 0.76 | 0.38 |

| Group 1 | Group 2 | Sample size | H | P-value | Q-value |
|---|---|---|---|---|---|
| Water sites (C, P, R) | | 45 | 3.9 | 0.14 | / |

| | | | | | |
|---|---|---|---|---|---|
| Pairwise comparisons | | | | | |
| C | P | 35 | 4.2 | 0.04 | 0.12 |
| C | R | 25 | 1.17 | 0.28 | 0.42 |
| P | R | 30 | 0.1 | 0.91 | 0.91 |

| Group 1 | Group 2 | Sample size | H | P-value | Q-value |
|---|---|---|---|---|---|
| Water sites (C2, C4, GU, OL, P1, P2) | | 45 | 4.84 | 0.43 | / |
| Pairwise comparisons | | | | | |
| C2 | C4 | 15 | 0.003 | 0.95 | 1 |
| C2 | GU | 12 | 1.64 | 0.2 | 0.65 |
| C2 | OL | 10 | 0.18 | 0.67 | 0.88 |
| C2 | P1 | 16 | 1.7 | 0.19 | 0.65 |
| C2 | P2 | 16 | 3.61 | 0.06 | 0.65 |
| C4 | GU | 15 | 1.26 | 0.26 | 0.65 |
| C4 | OL | 13 | 0 | 1 | 1 |
| C4 | P1 | 19 | 1.3 | 0.25 | 0.65 |
| C4 | P2 | 19 | 2.17 | 0.14 | 0.65 |
| GU | OL | 10 | 0.72 | 0.39 | 0.81 |
| GU | P1 | 16 | 0.1 | 0.74 | 0.88 |
| GU | P2 | 16 | 0.19 | 0.66 | 0.88 |
| OL | P1 | 14 | 0.6 | 0.43 | 0.82 |
| OL | P2 | 14 | 0.32 | 0.57 | 0.88 |
| P1 | P2 | 20 | 0.09 | 0.76 | 0.88 |

### trnL - Discarding Chlorophyta

| Group 1 | Group 2 | Sample size | H | P-value |
|---|---|---|---|---|
| Air | Water | 75 | 38.07 | **<0.001** |
| S1 | S2 | 30 | 0.34 | 0.56 |

| Group 1 | Group 2 | Sample size | H | P-value | Q-value |
|---|---|---|---|---|---|
| Water sites (C, P, R) | | 45 | 4.8 | 0.09 | / |

| Pairwise comparisons | | | | | |
|---|---|---|---|---|---|
| C | P | 35 | 4.3 | 0.04 | 0.11 |
| C | R | 25 | 2.3 | 0.12 | 0.19 |
| P | R | 30 | 0.23 | 0.63 | 0.63 |

| Group 1 | Group 2 | Sample size | H | P-value | Q-value |
|---|---|---|---|---|---|
| Water sites (C2, C4, GU, OL, P1, P2) | | 45 | 5.6 | 0.35 | / |
| Pairwise comparisons | | | | | |
| C2 | C4 | 15 | 1.12 | 0.29 | 0.75 |
| C2 | GU | 12 | 0.64 | 0.42 | 0.86 |
| C2 | OL | 10 | 0.18 | 0.67 | 0.91 |
| C2 | P1 | 16 | 1.06 | 0.3 | 0.75 |
| C2 | P2 | 16 | 1.06 | 0.3 | 0.75 |
| C4 | GU | 15 | 3.57 | 0.058 | 0.44 |
| C4 | OL | 13 | 0.48 | 0.48 | 0.86 |
| C4 | P1 | 19 | 3.84 | 0.05 | 0.44 |
| C4 | P2 | 19 | 2.41 | 0.12 | 0.6 |
| GU | OL | 10 | 0.1 | 0.75 | 0.91 |
| GU | P1 | 16 | 0.01 | 0.91 | 0.91 |
| GU | P2 | 16 | 0.047 | 0.83 | 0.91 |
| OL | P1 | 14 | 0.32 | 0.57 | 0.86 |
| OL | P2 | 14 | 0.32 | 0.57 | 0.86 |
| P1 | P2 | 20 | 0.01 | 0.9 | 0.91 |

**S5b. PERMANOVA results**

PERMANOVA analysis was performed for each marker gene and for a subgroup of taxa considering the type of medium, the macro category site and the sampling site. In particular, macro categories are P (P1 e P2) , C (C2 e C4) and R (Guisa = GU e Olona = OL). Analysis was carried out considering Jaccard distance and 999 permutations.
Below the results for each comparison and marker gene.

**V9 18S**

| Group 1 | Group 2 | Sample size | Pseudo-F | P-value |
|---|---|---|---|---|
| Air | Water | 81 | 10.76 | **0.001** |
| S1 | S2 | 34 | 0.93 | 0.621 |

| Group 1 | Group 2 | Sample size | Pseudo-F | P-value | Q-value |
|---|---|---|---|---|---|
| Water sites (C, P, R) | | 47 | 5.09 | **0.001** | / |
| Pairwise comparisons | | | | | |
| C | P | 37 | 6.02 | 0.001 | **0.001** |
| C | R | 26 | 4.17 | 0.001 | **0.001** |
| P | R | 31 | 4.75 | 0.001 | **0.001** |

| Group 1 | Group 2 | Sample size | Pseudo-F | P-value | Q-value |
|---|---|---|---|---|---|
| Water sites (C2, C4, GU, OL, P1, P2) | | 47 | 3.02 | **0.001** | / |
| Pairwise comparisons | | | | | |
| C2 | C4 | 16 | 1.38 | 0.139 | **0.139** |
| C2 | GU | 13 | 2.96 | 0.001 | **0.001** |
| C2 | OL | 11 | 2.34 | 0.007 | **0.008** |
| C2 | P1 | 17 | 3.3 | 0.001 | **0.001** |
| C2 | P2 | 18 | 3.5 | 0.001 | **0.001** |
| C4 | GU | 15 | 3.35 | 0.001 | **0.001** |
| C4 | OL | 13 | 2.57 | 0.002 | **0.003** |
| C4 | P1 | 19 | 4.18 | 0.001 | **0.001** |
| C4 | P2 | 20 | 4.52 | 0.001 | **0.001** |
| GU | OL | 10 | 1.34 | 0.035 | **0.037** |
| GU | P1 | 16 | 3.5 | 0.001 | **0.001** |
| GU | P2 | 17 | 3.75 | 0.001 | **0.001** |
| OL | P1 | 14 | 2.64 | 0.001 | **0.001** |
| OL | P2 | 15 | 2.87 | 0.001 | **0.001** |
| P1 | P2 | 21 | 1.82 | 0.005 | **0.006** |

**V9 18S - Eukaryota**

| Group 1 | Group 2 | Sample size | Pseudo-F | P-value |
|---|---|---|---|---|
| Air | Water | 81 | 11.05 | **0.001** |
| S1 | S2 | 34 | 0.94 | 0.58 |

| Group 1 | Group 2 | Sample size | Pseudo-F | P-value | Q-value |
|---|---|---|---|---|---|
| Water sites (C, P, R) | | 47 | 4.89 | **0.001** | / |
| Pairwise comparisons | | | | | |
| C | P | 37 | 5.62 | 0.001 | **0.001** |
| C | R | 26 | 4.06 | 0.001 | **0.001** |
| P | R | 31 | 4.70 | 0.001 | **0.001** |

| Group 1 | Group 2 | Sample size | Pseudo-F | P-value | Q-value |
|---|---|---|---|---|---|
| Water sites (C2, C4, GU, OL, P1, P2) | | 47 | 2.93 | **0.001** | / |
| Pairwise comparisons | | | | | |
| C2 | C4 | 16 | 1.37 | 0.14 | 0.14 |
| C2 | GU | 13 | 2.87 | 0.001 | **0.002** |
| C2 | OL | 11 | 2.31 | 0.012 | **0.014** |
| C2 | P1 | 17 | 3.13 | 0.002 | **0.002** |
| C2 | P2 | 18 | 3.36 | 0.001 | **0.002** |
| C4 | GU | 15 | 3.20 | 0.001 | **0.002** |
| C4 | OL | 13 | 2.52 | 0.002 | **0.003** |
| C4 | P1 | 19 | 3.93 | 0.001 | **0.002** |
| C4 | P2 | 20 | 4.27 | 0.001 | **0.002** |
| GU | OL | 10 | 1.3 | 0.043 | **0.046** |
| GU | P1 | 16 | 3.44 | 0.002 | **0.002** |
| GU | P2 | 17 | 3.70 | 0.001 | **0.002** |
| OL | P1 | 14 | 2.63 | 0.002 | **0.002** |
| OL | P2 | 15 | 2.88 | 0.002 | **0.002** |
| P1 | P2 | 21 | 1.87 | 0.006 | **0.007** |

**V9 18S - Metazoa**

| Group 1 | Group 2 | Sample size | Pseudo-F | P-value |
|---|---|---|---|---|
| Air | Water | 61 | 4.46 | **0.001** |
| S1 | S2 | 27 | 1.08 | 0.25 |

| Group 1 | Group 2 | Sample size | Pseudo-F | P-value | Q-value |
|---|---|---|---|---|---|
| Water sites (C, P, R) | | 44 | 3.28 | **0.001** | / |
| Pairwise comparisons | | | | | |
| C | P | 37 | 4.48 | 0.001 | **0.001** |
| C | R | 23 | 2.31 | 0.001 | **0.001** |
| P | R | 28 | 2.52 | 0.001 | **0.001** |

| Group 1 | Group 2 | Sample size | Pseudo-F | P-value | Q-value |
|---|---|---|---|---|---|
| Water sites (C2, C4, GU, OL, P1, P2) | | 44 | 2.35 | **0.001** | / |
| Pairwise comparisons | | | | | |
| C2 | C4 | 16 | 1.63 | 0.02 | **0.02** |
| C2 | GU | 10 | 1.68 | 0.03 | **0.03** |
| C2 | OL | 11 | 1.86 | 0.02 | **0.02** |
| C2 | P1 | 17 | 2.77 | 0.001 | **0.002** |
| C2 | P2 | 18 | 2.94 | 0.001 | **0.002** |
| C4 | GU | 12 | 1.75 | 0.004 | **0.006** |
| C4 | OL | 13 | 1.9 | 0.003 | **0.006** |
| C4 | P1 | 19 | 3.47 | 0.001 | **0.002** |
| C4 | P2 | 20 | 3.92 | 0.001 | **0.002** |
| GU | OL | 7 | 0.95 | 0.48 | 0.48 |
| GU | P1 | 13 | 1.76 | 0.006 | **0.009** |
| GU | P2 | 14 | 1.97 | 0.01 | **0.013** |
| OL | P1 | 14 | 2.02 | 0.004 | **0.006** |
| OL | P2 | 15 | 2.14 | 0.001 | **0.002** |
| P1 | P2 | 21 | 2.24 | 0.001 | **0.002** |

### ITS2

| Group 1 | Group 2 | Sample size | Pseudo-F | P-value |
|---|---|---|---|---|
| Air | Water | 62 | 6.39 | **0.001** |
| S1 | S2 | 22 | 0.77 | 0.92 |

| Group 1 | Group 2 | Sample size | Pseudo-F | P-value | Q-value |
|---|---|---|---|---|---|
| Water sites (C, P, R) | | 40 | 3.59 | **0.001** | / |
| Pairwise comparisons | | | | | |
| C | P | 31 | 4.59 | 0.001 | **0.001** |
| C | R | 22 | 3.24 | 0.001 | **0.001** |
| P | R | 27 | 2.8 | 0.001 | **0.001** |

| Group 1 | Group 2 | Sample size | Pseudo-F | P-value | Q-value |
|---|---|---|---|---|---|
| Water sites (C2, C4, GU, OL, P1, P2) | | 40 | 2.23 | **0.01** | / |
| Pairwise comparisons | | | | | |
| C2 | C4 | 13 | 1.00 | 0.445 | 0.445 |
| C2 | GU | 10 | 2.11 | 0.006 | **0.009** |
| C2 | OL | 9 | 1.79 | 0.012 | **0.015** |
| C2 | P1 | 13 | 2.16 | 0.009 | **0.012** |
| C2 | P2 | 15 | 2.58 | 0.001 | **0.003** |
| C4 | GU | 13 | 2.78 | 0.001 | **0.003** |
| C4 | OL | 12 | 2.2 | 0.004 | **0.007** |
| C4 | P1 | 16 | 3.11 | 0.002 | **0.004** |
| C4 | P2 | 18 | 3.63 | 0.001 | **0.003** |
| GU | OL | 9 | 1.25 | 0.06 | 0.07 |
| GU | P1 | 13 | 2.32 | 0.001 | **0.003** |
| GU | P2 | 15 | 2.41 | 0.001 | **0.003** |
| OL | P1 | 12 | 1.83 | 0.006 | **0.009** |
| OL | P2 | 14 | 1.86 | 0.002 | **0.004** |
| P1 | P2 | 18 | 1.49 | 0.047 | 0.054 |

**ITS2 - Fungi**

| Group 1 | Group 2 | Sample size | Pseudo-F | P-value |
|---|---|---|---|---|
| Air | Water | 35 | 3.13 | **0.001** |
| S1 | S2 | 22 | 0.74 | 0.93 |

| Group 1 | Group 2 | Sample size | Pseudo-F | P-value | Q-value |
|---|---|---|---|---|---|
| Water sites (C, P, R) | | 35 | 2.03 | **0.001** | / |
| Pairwise comparisons | | | | | |
| C | P | 26 | 1.59 | 0.004 | **0.004** |
| C | R | 17 | 1.95 | 0.002 | **0.003** |
| P | R | 27 | 2.5 | 0.001 | **0.003** |

| Group 1 | Group 2 | Sample size | Pseudo-F | P-value | Q-value |
|---|---|---|---|---|---|
| Water sites (C2, C4, GU, OL, P1, P2) | | 35 | 1.56 | **0.001** | / |
| Pairwise comparisons | | | | | |
| C2 | C4 | 8 | 0.9 | 0.62 | 0.62 |
| C2 | GU | 10 | 2.05 | 0.007 | **0.021** |
| C2 | OL | 9 | 1.32 | 0.11 | 0.13 |
| C2 | P1 | 13 | 1.4 | 0.039 | 0.053 |
| C2 | P2 | 15 | 1.6 | 0.006 | **0.021** |
| C4 | GU | 8 | 1.7 | 0.018 | **0.038** |
| C4 | OL | 7 | 1.1 | 0.037 | 0.053 |
| C4 | P1 | 11 | 1.21 | 0.28 | 0.28 |
| C4 | P2 | 13 | 1.2 | 0.1 | 0.13 |
| GU | OL | 9 | 1.37 | 0.026 | **0.043** |
| GU | P1 | 13 | 2.4 | 0.22 | **0.015** |
| GU | P2 | 15 | 2.35 | 0.001 | **0.015** |
| OL | P1 | 12 | 1.6 | 0.009 | **0.022** |
| OL | P2 | 14 | 1.51 | 0.003 | **0.015** |

| | | | | | |
|---|---|---|---|---|---|
| P1 | P2 | 18 | 1.44 | 0.023 | 0.043 |

**trnL**

| Group 1 | Group 2 | Sample size | Pseudo-F | P-value |
|---|---|---|---|---|
| Air | Water | 73 | 5.65 | 0.001 |
| S1 | S2 | 30 | 1.18 | 0.063 |

| Group 1 | Group 2 | Sample size | Pseudo-F | P-value | Q-value |
|---|---|---|---|---|---|
| Water sites (C, P, R) | | 40 | 3.15 | 0.001 | / |
| Pairwise comparisons | | | | | |
| C | P | 30 | 4.36 | 0.001 | 0.001 |
| C | R | 25 | 3.15 | 0.001 | 0.001 |
| P | R | 25 | 2.95 | 0.001 | 0.001 |

| Group 1 | Group 2 | Sample size | Pseudo-F | P-value | Q-value |
|---|---|---|---|---|---|
| Water sites (C2, C4, GU, OL, P1, P2) | | 40 | 2.24 | 0.001 | / |
| Pairwise comparisons | | | | | |
| C2 | C4 | 15 | 1.11 | 0.26 | 0.26 |
| C2 | GU | 12 | 2.12 | 0.002 | 0.003 |
| C2 | OL | 10 | 2.11 | 0.005 | 0.006 |
| C2 | P1 | 12 | 2.59 | 0.002 | 0.003 |
| C2 | P2 | 15 | 2.58 | 0.001 | 0.003 |
| C4 | GU | 15 | 2.42 | 0.001 | 0.003 |
| C4 | OL | 13 | 2.31 | 0.002 | 0.003 |
| C4 | P1 | 15 | 3.22 | 0.001 | 0.003 |
| C4 | P2 | 18 | 3.22 | 0.001 | 0.003 |
| GU | OL | 10 | 1.27 | 0.008 | 0.009 |
| GU | P1 | 12 | 2.38 | 0.001 | 0.003 |
| GU | P2 | 15 | 2.5 | 0.001 | 0.003 |

| | | | | | |
|---|---|---|---|---|---|
| OL | P1 | 10 | 1.91 | 0.003 | **0.004** |
| OL | P2 | 13 | 2.03 | 0.002 | **0.003** |
| P1 | P2 | 15 | 1.51 | 0.029 | **0.031** |

**trnL - Streptophyta**

| Group 1 | Group 2 | Sample size | Pseudo-F | P-value |
|---|---|---|---|---|
| Air | Water | 75 | 5.21 | **0.001** |
| S1 | S2 | 30 | 1.15 | 0.12 |

| Group 1 | Group 2 | Sample size | Pseudo-F | P-value | Q-value |
|---|---|---|---|---|---|
| Water sites (C, P, R) | | 45 | 3.54 | **0.001** | / |
| Pairwise comparisons | | | | | |
| C | P | 35 | 4.21 | 0.001 | **0.001** |
| C | R | 25 | 3.12 | 0.001 | **0.001** |
| P | R | 30 | 3.17 | 0.001 | **0.001** |

| Group 1 | Group 2 | Sample size | Pseudo-F | P-value | Q-value |
|---|---|---|---|---|---|
| Water sites (C2, C4, GU, OL, P1, P2) | | 45 | 2.3 | **0.001** | / |
| Pairwise comparisons | | | | | |
| C2 | C4 | 15 | 1.09 | 0.26 | 0.26 |
| C2 | GU | 12 | 2 | 0.001 | **0.002** |
| C2 | OL | 10 | 2 | 0.006 | **0.007** |
| C2 | P1 | 16 | 2.51 | 0.001 | **0.002** |
| C2 | P2 | 16 | 2.3 | 0.001 | **0.002** |
| C4 | GU | 15 | 2.41 | 0.001 | **0.007** |
| C4 | OL | 13 | 2.3 | 0.004 | **0.005** |
| C4 | P1 | 19 | 3.41 | 0.001 | **0.007** |
| C4 | P2 | 19 | 3.08 | 0.001 | **0.007** |
| GU | OL | 10 | 1.19 | 0.037 | **0.039** |
| GU | P1 | 15 | 2.72 | 0.001 | **0.007** |

| | | | | | |
|---|---|---|---|---|---|
| GU | P2 | 16 | 2.56 | 0.001 | **0.007** |
| OL | P1 | 14 | 2.06 | 0.001 | **0.007** |
| OL | P2 | 14 | 2 | 0.001 | **0.007** |
| P1 | P2 | 20 | 1.79 | 0.004 | **0.005** |

**trnL - Discarding Chlorophyta**

| Group 1 | Group 2 | Sample size | Pseudo-F | P-value |
|---|---|---|---|---|
| Air | Water | 75 | 5.23 | **0.001** |
| S1 | S2 | 30 | 1.62 | 0.086 |

| Group 1 | Group 2 | Sample size | Pseudo-F | P-value | Q-value |
|---|---|---|---|---|---|
| Water sites (C, P, R) | | 45 | 3.55 | **0.001** | / |
| Pairwise comparisons | | | | | |
| C | P | 35 | 4.42 | 0.001 | **0.001** |
| C | R | 25 | 3.07 | 0.001 | **0.001** |
| P | R | 30 | 3 | 0.001 | **0.001** |

| Group 1 | Group 2 | Sample size | Pseudo-F | P-value | Q-value |
|---|---|---|---|---|---|
| Water sites (C2, C4, GU, OL, P1, P2) | | 45 | 2.26 | **0.001** | / |
| Pairwise comparisons | | | | | |
| C2 | C4 | 15 | 1.06 | 0.35 | 0.35 |
| C2 | GU | 12 | 2.02 | 0.001 | **0.002** |
| C2 | OL | 10 | 2 | 0.006 | **0.007** |
| C2 | P1 | 16 | 2.66 | 0.001 | **0.002** |
| C2 | P2 | 16 | 2.34 | 0.001 | **0.002** |
| C4 | GU | 15 | 2.40 | 0.001 | **0.002** |
| C4 | OL | 13 | 2.23 | 0.002 | **0.003** |
| C4 | P1 | 19 | 3.44 | 0.001 | **0.002** |
| C4 | P2 | 19 | 3.06 | 0.001 | **0.002** |
| GU | OL | 10 | 1.24 | 0.005 | **0.006** |

| | | | | | |
|---|---|---|---|---|---|
| GU | P1 | 16 | 2.57 | 0.001 | 0.002 |
| GU | P2 | 16 | 2.42 | 0.001 | 0.002 |
| OL | P1 | 14 | 2.02 | 0.001 | 0.002 |
| OL | P2 | 14 | 1.94 | 0.002 | 0.003 |
| P1 | P2 | 20 | 1.64 | 0.009 | 0.01 |

### S6. PCoA Analysis considering Time and Site metadata

PCoa calculated based on Jaccard metric, considering features.
In the legend: macro category site and month of sampling.

### V9 18S

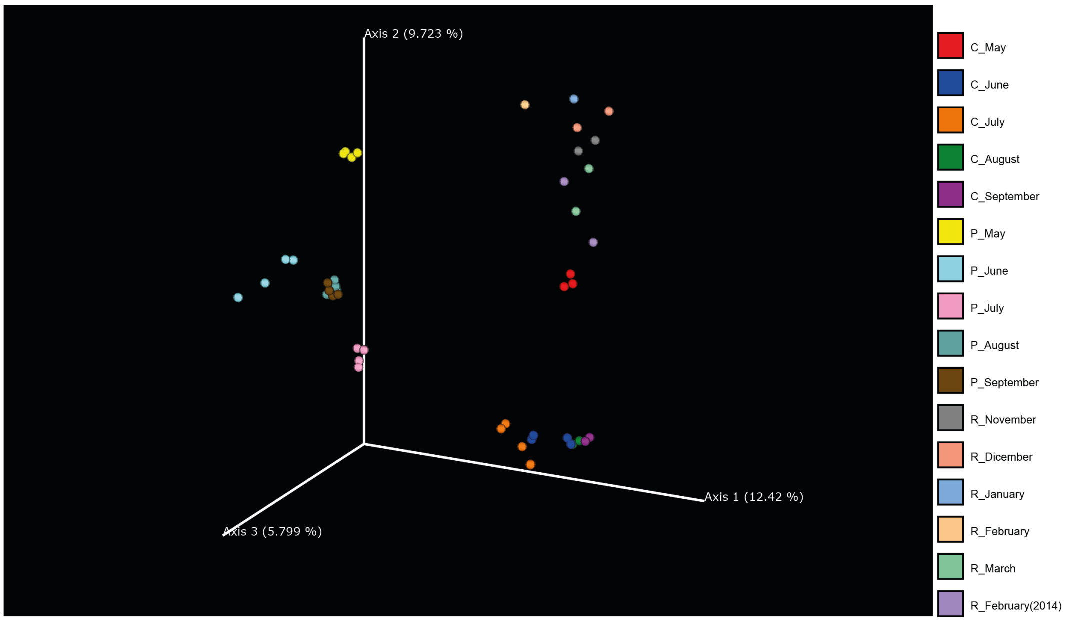

**ITS2**

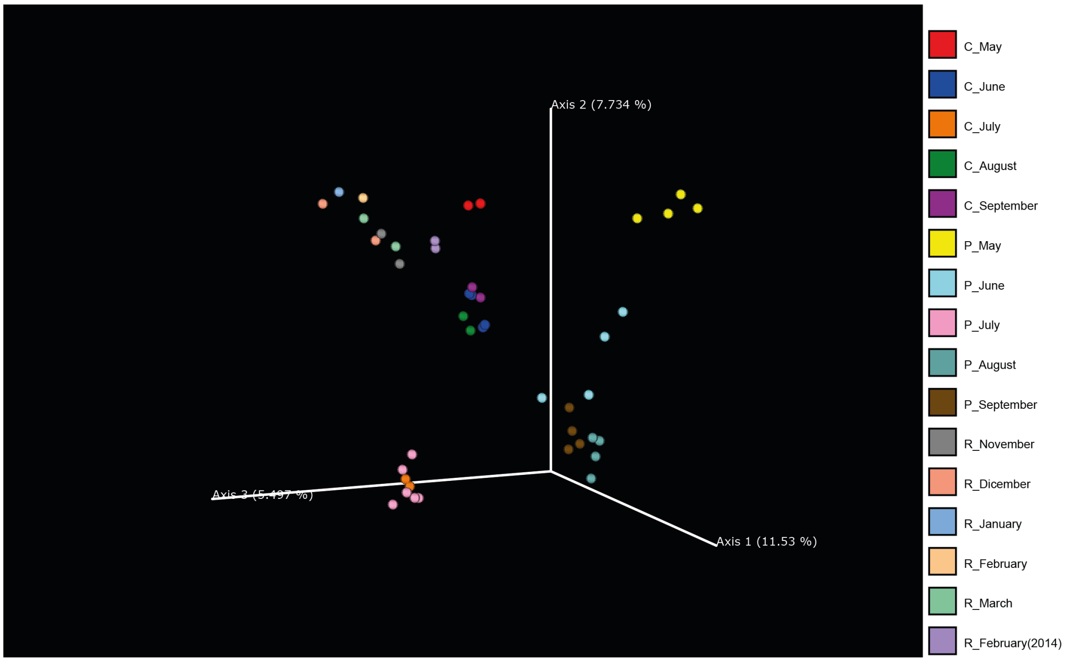

Axis 2 (7.734 %)
Axis 3 (5.497 %)
Axis 1 (11.53 %)
C_May
C_June
C_July
C_August
C_September
P_May
P_June
P_July
P_August
P_September
R_November
R_Dicember
R_January
R_February
R_March
R_February(2014)


**trnL**

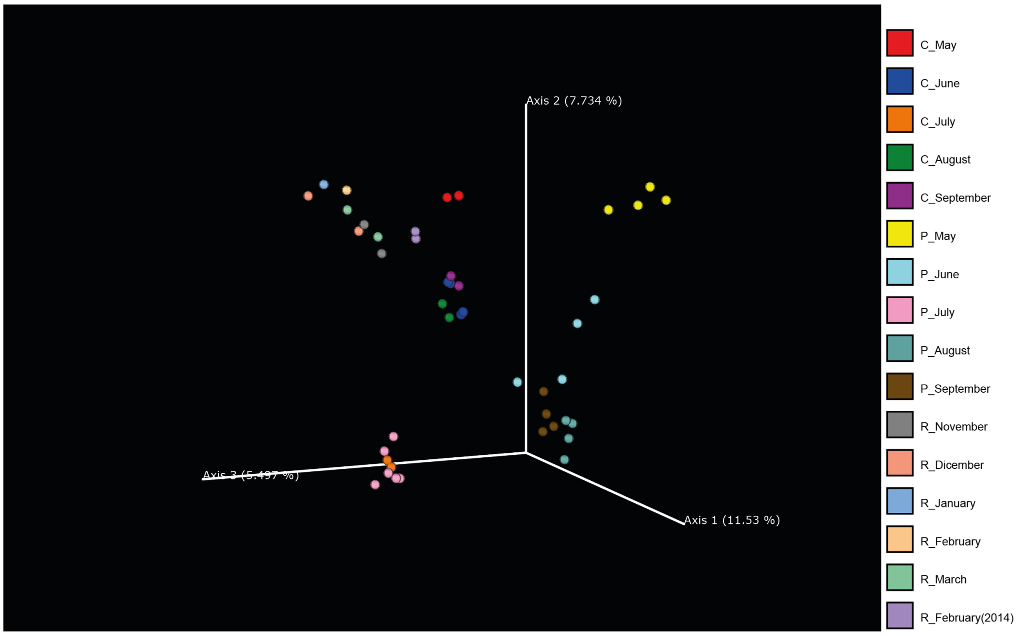

Axis 2 (7.734 %)
Axis 3 (5.497 %)
Axis 1 (11.53 %)
C_May
C_June
C_July
C_August
C_September
P_May
P_June
P_July
P_August
P_September
R_November
R_Dicember
R_January
R_February
R_March
R_February(2014)

#### S7. Interactive sunburst files

Interactive sunburst charts are provided in order to better explore the composition of the taxonomy assignment of the three markers (HTML format).
In the Supplementary directory there are

- S7a: sunburst taxa v9;
- S7b: sunburst taxa ITS2;
- S7c: sunburst taxa trnL.

#### S8. Feature table

The feature tables of the three markers were merged to obtain a unique feature table.
File: S8 - feature table (TSV format).

#### S9. Taxonomy tables

For each marker, taxonomy assignments can be found in the Supplementary directory.
Files (TSV format):

- S9a: 18S v9 taxonomy assignments;
- S9b: ITS2 taxonomy assignments;
- S9c: trnL taxonomy assignments.

**S10. ML sample classifier**

**ITS2**

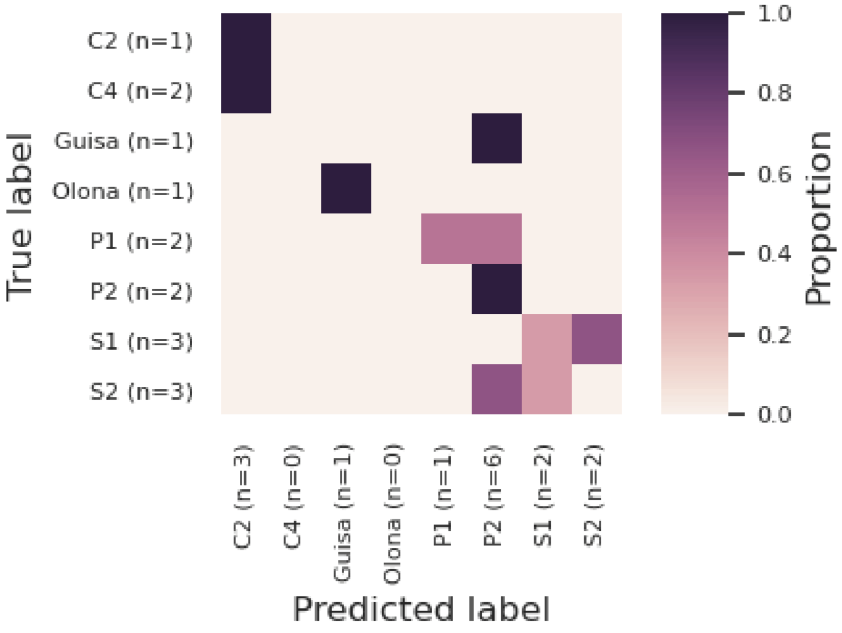


a)

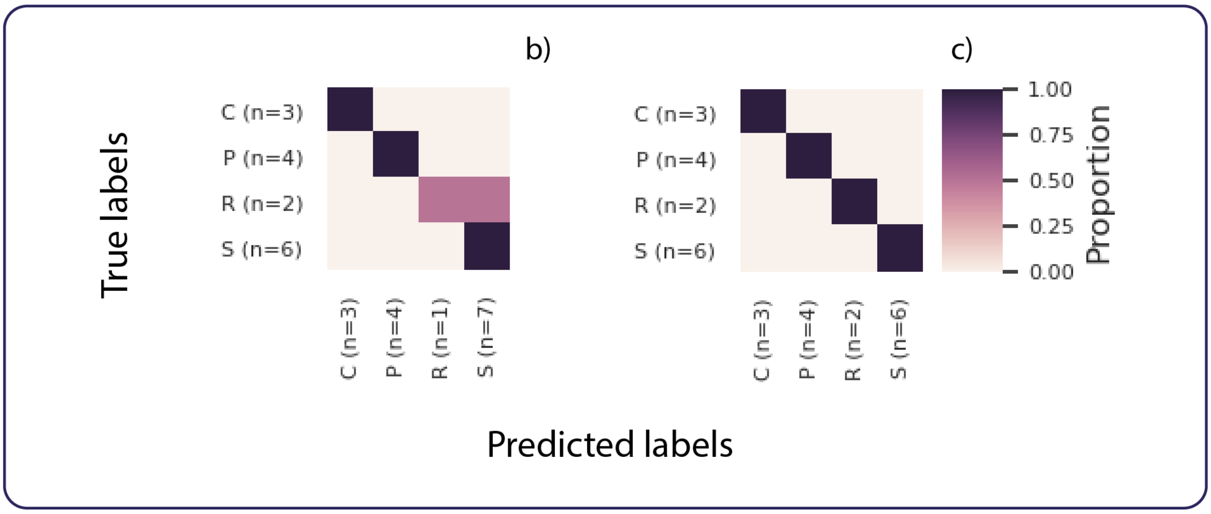


Machine learning of ITS marker: results were reported considering a) sequences per sites and considering the division of sites into macro categories of b) sequences and c) Fungi.

**trnL**

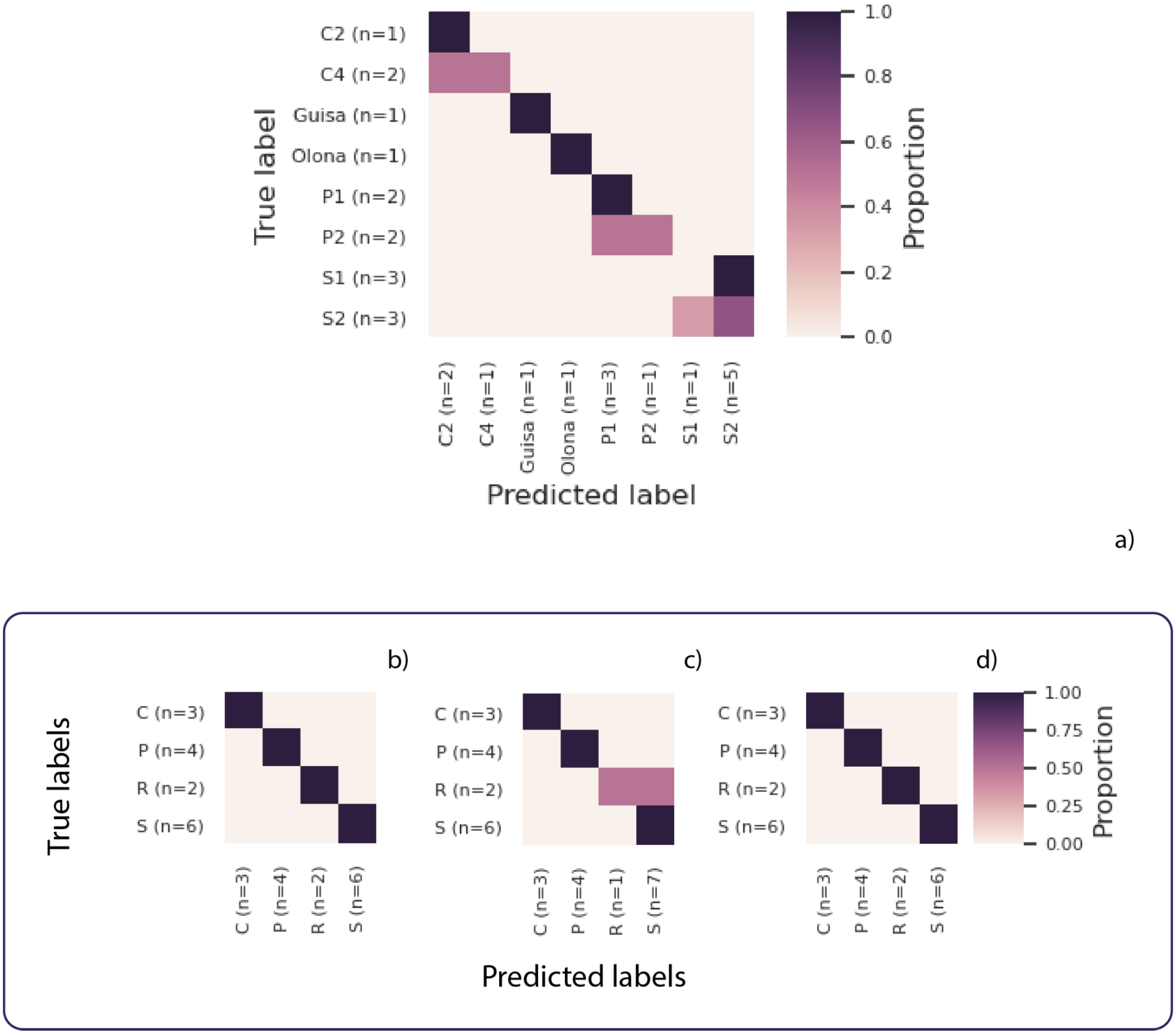


Machine learning of trnL marker: results were reported considering a) sequences per sites and considering the division of sites into macro categories of b) sequences, c) Streptophyta, d) excluding Chlorophyta.
